# Threshold accumulation of a constitutive protein explains *E. coli* cell division behavior in nutrient upshifts

**DOI:** 10.1101/2020.08.03.233908

**Authors:** Mia Panlilio, Jacopo Grilli, Giorgio Tallarico, Ilaria Iuliani, Bianca Sclavi, Pietro Cicuta, Marco Cosentino Lagomarsino

**Affiliations:** Cavendish Laboratory, Cambridge University, Cambridge, United Kingdom; The Abdus Salam International Centre for Theoretical Physics (ICTP), Strada Costiera 11, 34014 Trieste, Italy; Dipartimento di Fisica, Università degli Studi di Milano, via Celoria 16 Milano, Italy; Sorbonne Université, Campus Pierre and Marie Curie, 4 Place Jussieu 75005, Paris, France; CNRS, UMR7238, 4 Place Jussieu 75005, Paris, France; IFOM Foundation, FIRC Institute of Molecular Oncology, Milan, Italy; Dipartimento di Fisica, Università degli Studi di Milano, and I.N.F.N, via Celoria 16 Milano, Italy

## Abstract

Despite of a boost of recent progress in dynamic single-cell measurements and analyses in *E. coli*, we still lack a mechanistic understanding of the determinants of the decision to divide. Specifically, the debate is open regarding the processes linking growth and chromosome replication to division, and on the molecular origin of the observed “adder correlations”, whereby cells divide adding roughly a constant volume independent of their initial volume. In order to gain insight into these questions, we interrogate dynamic size-growth behavior of single cells across nutrient upshifts with a high-precision microfluidic device. We find that the division rate changes quickly after nutrients change, much before growth rate goes to a steady state, and in a way that adder correlations are robustly conserved. Comparison of these data to simple mathematical models falsifies proposed mechanisms where replication-segregation or septum completion are the limiting step for cell division. Instead, we show that the accumulation of a putative constitutively expressed “P-sector divisor” protein explains the behavior during the shift.

**Significance statement:** The mechanism leading to cell division in the bacterium *E. coli* is unknown, but we know that it results in adding a roughly constant size every cell cycle, regardless of size at birth. While most available studies try to infer information on cell division from steadily dividing cells in constant nutrient conditions, this study leverages on a high-resolution device to monitor single-cell growth division upon nutrient changes. Comparing these data with different mathematical models, the authors are able to discriminate among fundamentally different mechanisms of cell division control, and they show that the data support a model where an unregulated protein accumulates to a threshold and triggers division.

## INTRODUCTION

To divide, a cell needs to coordinate the allocation and duration of multiple processes, including metabolism, maintenance of cellular compartments and faithful replication and segregation of chromosomes [1–5]. Hence, an accurately timed decision to divide may have primary importance for survival and fitness. Achieving this accuracy likely requires a combination of upstream scheduling and downstream control, many aspects of which are only partially known [2, 6–8].

Classic work has addressed cell size control based on population measurements [5], drawing conclusions that are increasingly challenged by recent findings [7, 9–12]. In particular, we do not know if there is one fixed rate-limiting process setting division, and in this case if it is related to chromosome synthesis and segregation, or to cell-surface related processes. Recent work on the behavior of single cells has identified clear phenomenological patterns by which cells decide to divide. These can be characterized as correlation patterns between size-growth variables such as added or multiplicative growth [13–17]. Nearly all of the available work has focused on steady-state (balanced growth) single-cell data, and established that cell size is not the only variable entering the decision to divide [18], and that cells follow a remarkably simple and robust pattern for setting division [13, 19–21], characterized by low or no correlations between the added size over one cell cycle and the size at birth (near- “adder” correlations). However, we still largely ignore both the time hierarchy (or more complex scheduling plan) and the molecular players at the basis of the observed near-adder behavior.

More generally, models founded on different mechanisms for division control can give rise to equivalent predictions in steady growth. Steady conditions only allow the exploration of correlations (or more generally joint distributions) between quantities involved in cell-size control [15]. For instance, the evidence of the adder behavior comes from the observation of a lack of correlation between added size and initial size at the single-cell level [20]. In order to discriminate between alternative mechanisms one needs to go beyond correlations and to explore causal relationships between the variables involved in size control. Exploring the dynamics of size control in non-steady conditions, e.g., during a nutrient shift, has the potential to allow one to disentangle alternative models based on different causal relationships.

Near-adder behavior, also observed between consecutive initiations [9, 22] may derive from accumulation and trigger of an “initiator” protein setting replication initiation [16, 23, 24]. The initiation of replication is known to be effected by a critical accumulation of ATP-bound DnaA. While some molecular mechanisms involved in DNA opening and replisome assembly have been identified, it is still not known how they contribute to setting the timing of initiation of DNA replication in different growth conditions or how they can contribute to cell-cycle progression in single cells [10, 25, 26]. In particular, additional players and processes such as the alarmone ppGpp and DNA supercoiling are implicated [27, 28], and DnaA itself might not even be rate-limiting for replication initiation in all growth conditions [29]. The further assumption that chromosome replication-segregation limits cell division would explain the observed near-adder correlations between divisions [16], but this is also subject of intense debate, as there is still no agreement on whether the initiation process itself may be rate-limiting for cell division [7–9, 16, 22, 30, 31]. Independently from the chromosome cycle, cell division (and its adder correlations) could be set by the accumulation of a putative “divisor” factor, possibly related to the FtsZ division ring [9, 32] or to the synthesis of the septum [31]. Finally, both the division and initiation processes could be scheduled upstream by yet unidentified central cell-cycle regulators [2, 10], and downstream control processes could condition cell division to the completion of a set of necessary processes [7, 8].

In order to shed more light on cell division dynamics, one needs to access conditions where nutrient-imposed growth rate and division rate are decoupled, and look at the behavior of single cells. To this end, we followed the growth-division dynamics of cells undergoing nutrient upshifts in a high-resolution microfluidic device. In this device, cells can be kept in steady-state growth, and go from minimal M9 medium with Glucose, to the same medium supplemented with Casaminoacids. In these shifts, the change of amino acid levels was shown to alter the amount of free ppGpp, affecting translation and transcription of ribosomal promoters [33]. However, the division rate dynamics, and the division control behavior of single cells remain to be characterized. Our results identify the time scales involved in the changes of growth- and division-related variables, and characterize the correlation patterns between cell size-growth and timing of cell division emanating from the mechanisms of division control. Second, we use the comparison data with mathematical models to support or and falsify different underlying mechanistic models, such as initiation-limited division, accumulation of a putative divisor protein, and slaving of changes in division rate to changes in growth rate.

## RESULTS

### Long-term single cell tracking through a nutritional shift

Robust environmental control was achieved by cultivating the *E. coli* cells in a ‘mother machine’ microfluidic device [34, 35]. Fresh growth medium was fed into the device and cells were observed through a timed switch from stringent (M9 + 0.4% glucose, doubling time = 57 *±* 26 min) to rich media (M9 + 0.4% glucose + 0.5% casamino acids, doubling time = 37 *±* 13 min). Cells must allocate energy resources towards the biosynthesis of amino acids while in stringent media. When this metabolic limitation is relieved by the introduction of casamino acids into the external environment, cells grow and divide more quickly [36]. Sample images from these respective ‘slow’ and ‘fast’ steady growth states are shown in Fig. 1A. Consistent with Schaechter’s growth laws [37] and Woldringh’s original experiments [38], the increase in mean exponential growth rate 〈*α*_inst_〉 between steady states (0.0137 *±* 0.0052 min*^−^*^1^ to 0.0203 *±* 0.0062 min*^−^*^1^, Fig. 1B) saw a requisite increase in mean birth volume *(V*_0_〉 (1.15 *±* 0.26 *µ*m^3^ to 1.53 *±* 0.36 *µ*m^3^, Fig. 1B).

**FIG. 1.**
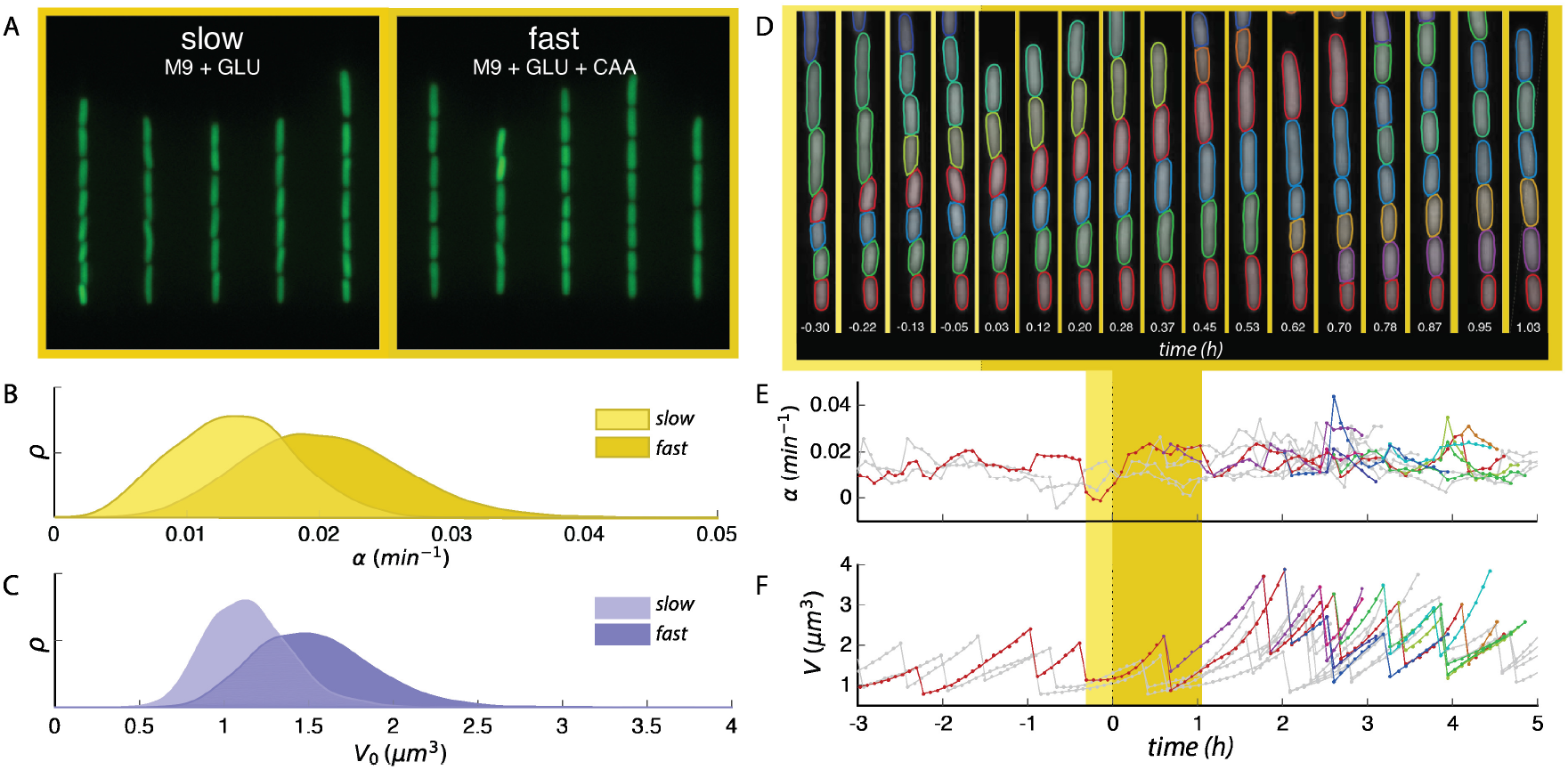
Robust and long term single cell tracking through a nutritional upshift. **A.** Snapshots of trapped cells in microfluidic device under stringent (left) and rich nutrient conditions (right) (false-coloured). **B.** Steady state distributions of growth rate *α_cc_* in the two growth media. **C.** Steady state distributions of birth volumes *V*_0_ for the sample populations in **B**. **D.** Sample segmentation and tracking before and after the growth media switch is implemented, time since switch shown in hours. **E.** Tracking the change of instantaneous growth rate following the switch for cells shown above. Yellow regions indicate time range spanned by segmented images, plotted line colour corresponds to the cell above with the same colour segmented boundary. **F.** Volume tracking of the same cell lines. Interdivision time *τ* identified as the interval between dramatic volume decreases.

The device configuration achieved high throughput without sacrificing spatio-temporal resolution, thereby enabling us to assess the system’s response to environmental perturbation at both population and single cell levels. Thousands of single cells were segmented and tracked for several generations before and after the nutritional shift Fig. 1D-F. Cell shape and size were measured from the segmented cell boundary (see Methods and SI Text) and used to determine individual growth rate and interdivision time. Specifically, the instantaneous growth rate of a single cell exhibiting exponential growth [34] is defined as 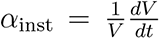 (Fig. 1E); cell division events can be found at the sudden and dramatic decreases in the volume of a tracked cell, yielding interdivision time *τ* (Fig. 1F).

### Non-adiabatic transition between steady growth states

In order to average equally and compare all the different cell-cycle quantities (interdivision time, growth rate from cycle-wide exponential fit, added volume), we associated their values to the time of division, then performed a sliding average. While different choices are possible, this is conservative with respect to quickly-changing variables, and, technically, it ensures that the sliding averages do not start changing before the actual time of the nutrient shift. For instantaneous quantities, such as as the growth rate and the production rates of the promoters, we also considered averages of the instantaneous (discrete) derivatives computed on single cells along lineages. These “instantaneous” averages are the appropriate choice in some instances, for example as input to the quantitative models.

Fig. 2 shows how the mean cell growth, cell division and cell size parameters follow a very complex dynamics across the nutrient upshift, characterized by multiple time scales and trends whereby the same quantity can both increase and decrease in different time windows.

**FIG. 2.**
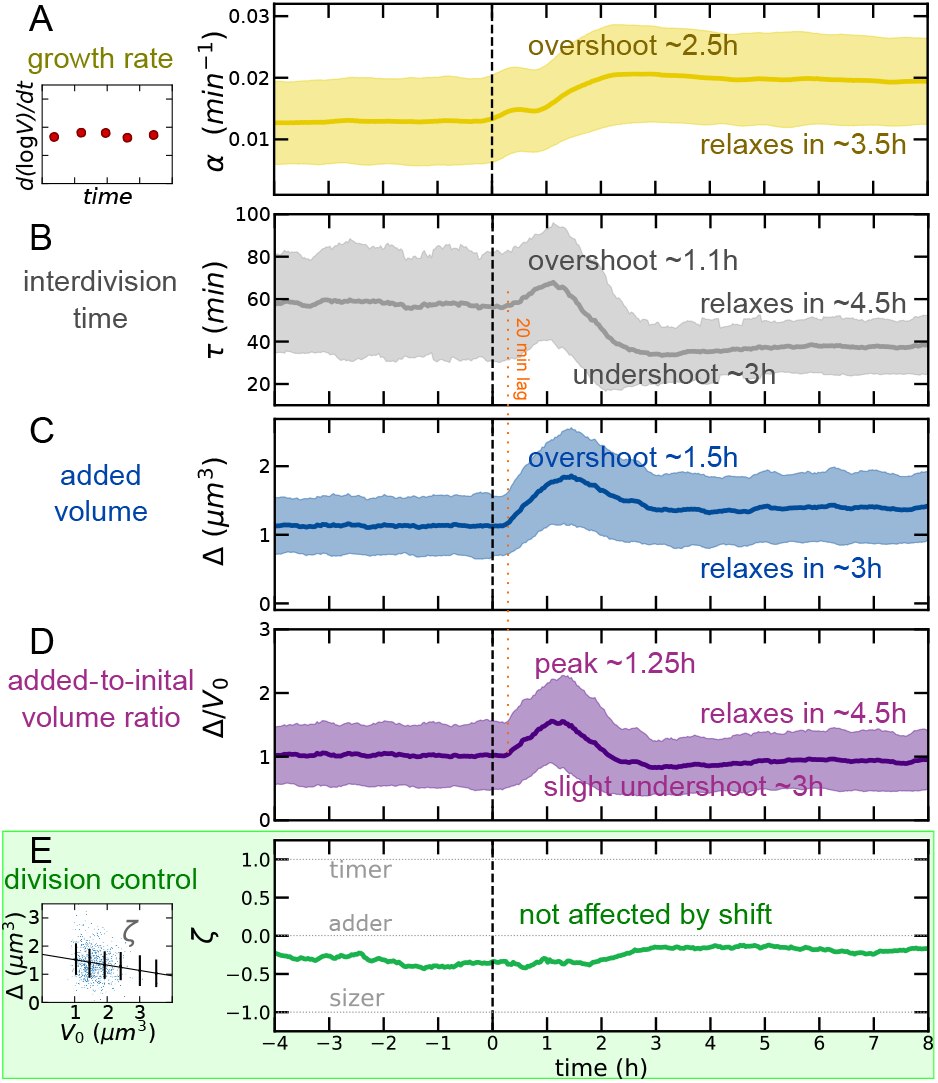
Near-adder behaviour is conserved through a nutrient upshift despite complex dynamics of growth and division processes. Media shift indicated at dotted vertical line. A) Mean growth rate (defined by a derivative of single-cell growth curves, as illustrated by the sketch on the left side) as a function of time in the experiment. Solid line indicates observable mean, and shaded regions indicate SD for all time series. B) Interdivision time transiently increases before reaching its new (lower) steady value across the upshift CD) Mean added volume shows an overshoot over more than 2 hours after the shift; the mean ratio of added to initial size is 1 at steady states. E) The division control shows near-adder behavior robustly across the shift. The adder plot (added size *vs* initial cell size, as illustrated by the sketch on the left side) quantifies division control by the slope *ζ* (−1 for sizer, 1 for timer, 0 for adder) [39].

Fig. 2A shows the trend of mean growth rate versus time in our data. Classically it is assumed that cells display ‘rate maintenance’ during an upshift, rapidly adopting the increased nutrient-imposed growth rate but continuing at the pre-shift interdivision time for about one generation [40, 41], indicating a divergence from the relation 〈*α*_inst_〈*τ*〉 = log 2. The combination of an increasing growth rate under constant interdivision time is thought by some to establish the increased volume characteristic of rich media [38, 42]. Indeed, some degree of deviation is necessary for cells to achieve the ultimately increased volume dictated by bacterial growth laws (see SI Text and ref.[10]). Here, we find behaviour far more complex than simple rate maintenance. In the minimal media, cells have a growth rate of 0.0138 *±* 0.0056 min*^−^*^1^. Upon upshift the growth rate increases gradually in agreement with early models of the upshift response [38] though initially it exceeds the new equlibrium value, reaching a maximum of 0.0219 *±* 0.0076 min*^−^*^1^ at *t* = 2.6 h before settling to 0.0204 *±* 0.0059 min*^−^*^1^.

Interdivision time exhibits a three-phase response (Fig. 2B). Initially increasing from its preshift value of 57 *±* 26 min to 67 *±* 28 min at *t* = 1.1 h, before quickly undershooting to a 34 *±* 14 min minimum two hours later. The equilibrium interdivision time is established after approximately 4.5 h in the nutrient-rich growth media to a value of 37 *±* 13 min (Fig. 2B). These non-complementary responses of growth and division processes can be summarized by the trajectory in the plane 〈*α*_inst_〉 vs. 1*/(τ)* “growth space” (SI Fig. S1A) we see that the nutritional upshift prompts the system to veer away from the “adiabatic” trajectory 〈*α*_inst_〉〈*τ*〉 = log 2 (dotted line), achieving a maximum of 0.968 at *t* = 1.3 h. That is, directly following the upshift, cells grow to a final volume that is on average up to 2.63 times their birth volume. The population 〈*α*_inst_〉〈*τ*〉 then decreases and actually undershoots log 2 at *t* = 3 h (green segment), indicating that cells slightly reduce their size transiently in relation to their mothers before reaching equilibrium. SI Fig. S1BCDE compare the trend of mean initial volume, cell width, growth rate and interdivision time. Cell width relaxes very slowly to the new condition [31]. Mean growth rate *(α)* and mean interdivision/doubling time *(τ)* at steady state are related through the equation 〈*α*_inst_〉〈*τ*〉 = log 2 [11, 43]. However, the data show clearly that the system does not obey this relation in the process of adapting to a different nutrient condition (SI Fig. S1F, we note also that in order to change mean size this relation needs to be transiently broken).

Finally, we can consider size changes across the shift. Simple calculations [13, 14, 20] show that at the steady state of balanced exponential growth, regardless of the mechanism of size control the average added size Δ and the average initial size have *V*_0_ to be equal in order to produce two daughter cells that are the same size as the mother, but the two quantities need to be decoupled outside of steady growth conditions, for example in environments with changing nutrient conditions. Following the nutritional upshift at *t* = 0, mean birth volume *V*_0_ and mean added volume Δ initially overshoot their increased equilibrium values in the new growth media (Fig. 2C and SI Fig. S1A). Specifically, birth volume peaks at roughly 111% of its new equilibrium (1.71 *±* 0.44 *µ*m^3^), reached at *t,..* 2.5 h and plateaus to 1.54 *±* 0.34 for *t ≥* 5 h (SI Fig. S1A). Similarly, added volume attains a maximum of 123% of its equilibrium value, reaching 2.00 *±* 0.78 *µ*m^3^ at *t,..* 1.5 h compared to its neq plateau value of 1.63 *±* 0.60 *µ*m^3^ reached for *t ≥* 4.5 h (SI Fig. S1). Growth rate varies slowly, but it immediately changes its trend after the shift. Instead, interdivision time and added volume appear to show a common delay of about 20 minutes (Fig. 2BCD). The delay is even longer (about 1h, SI Fig. S1) for initial volume[44]. The mean ratio of added to mean volume needs to be unitary at stationarity, and this is verified in the data (Fig. 2D). During the transition, this quantity follows a fairly symmetric peak to about 1.5 times its value in about 2.5h.

### Near-adder behaviour is conserved during the upshift

Recent studies have demonstrated the emergence of near-adder behaviour in bacterial cells, but the underlying biophysical and molecular mechanisms remain elusive [13, 20, 30, 43]. The size control strategy employed to adjust to a new growth environment is similarly mysterious since the existing adder observations have largely been confined to steady state conditions. Our data provide the opportunity to extend the analysis of the robustness and the determinants of adder behavior out of steady-state growth.

Adder size control can be tested by plotting the added size against the initial size of dividing cells [39], and evaluating the slope. We define this parameter as *ζ*. When *ζ* = 0 one has a perfect adder, i.e. cells add a size-independent volume to their initial size. The limits *ζ* = 1 and *ζ* = *−*1 correspond to a timer (no control) and a sizer (absolute size threshold Fig. 2E shows a strikingly robust trend of the measured near-adder behavior across the shift, despite of all the different time-scales involved in the geometry changes. To further characterize the adopted size control strategy and assess the contributions of the growth rate and interdivision time dynamics towards the overall size control, we also deployed an analysis developed in previous studies [15, 17] (see SI Fig. S2 and SI Text), whereby these contributions are inferred from the single-cell correlation patterns. This analysis confirms a near-adder behaviour that is uninterrupted by the environmental shift, effected by complementary dynamics between the timing- and growth-related components of size control.

### Nutrient-shift data are qualitatively incompatible with some proposed models for cell-division control

We first tested whether some previously proposed models could reproduce the complex behavior of division-related variables that we observed across the shift. To this end, we considered different models available in the literature (Fig 3A). Specifically, we ran single-cell simulations of the models proposed by Harris and Theriot [31] (“relative-rates” model), Ho and Amir [16] (“incremental” model), and the classic idea of “initiation sizer” (which we implemented here with size-uncoupled C+D period, see [7]), using both modeled (see methods) and sampled growth rate distributions across the shift (in the latter simulations each cell was assigned an exponential growth rate extracted from the measured ones in each particular time bin).

**FIG. 3.**
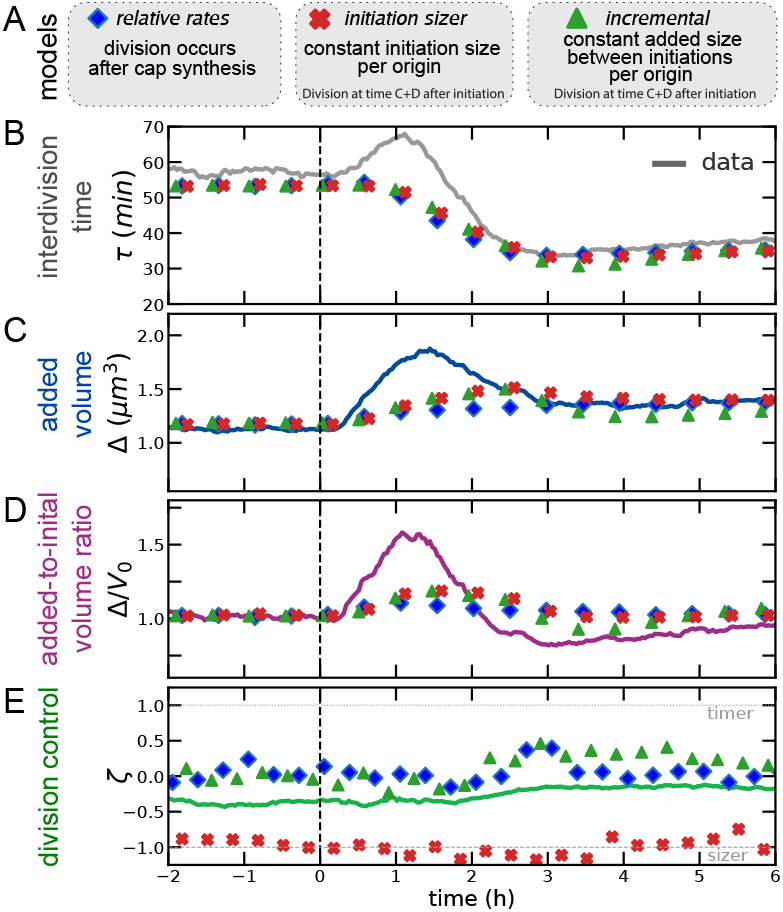
The nonsteady data across the nutrient shift falsify commonly assumed models for division dynamics. A) Summary of the models tested. BCD) Dynamics of interdivision time (B) added size and (C) across the shift cannot be recreated by existing size control models. In particular, all the tested models are too slow in reproducing the added-size dynamics and do not reproduce the initial increase in interdivision times. (D) The behavior of division control across the shift is steady across the shift for all three models considered.

The models are described in more detail in the methods section (see also SI text). The relative rates model assumes that chromosome replication-segregation is never limiting for cell division [9, 12], and that completion of synthesis of cap material (produced at the rate of surface synthesis) triggers division. A variant of the model may entail that target added size is proportional to width squared [31], and similar models assume that ring synthesis is limiting [32], but such variants are not crucial for our scopes. Both the incremental model and the initiation sizer model assume that replication-segregation are limiting for cell division. We did not consider the more complex scenario of concurrent time scales [7, 8] because of the unknown extra parameters used in this framework. The incremental model assumes that the chromosome is always limiting for cell division, and an inter-initiation adder (per origin) based on the cell current growth rate. By contrast, the initiation sizer model assumes a critical size per origin at initiation.

None of the considered envelope- or chromosome-limited models for cell division predict correctly the division time dynamics (Fig 3B). Specifically, no model predicts that the interdivision time in the data initially *increases*, before decreasing to its new steady state. Considering inter-division timing dynamics across the shift, under increasing *α*_inst_, in these models inter-division time can only decrease upon increase of growth rate, hence the models cannot reproduce the over/undershoot we observe for inter-division timing. Considering the dynamics of added size and initial size (Fig 3CD) all models show a delayed dynamics compared to the experimental data, and cannot predict the observed early overshoot in added volume. Buke and coworkers observe that the added size responds abruptly to changes in the ppGpp level, aided by transiently accelerated divisions upon a downshift, while growth adjusts on longer time scales [45]. Confirming this observation, in our upshift data the (counter-intuitive) initial increase of inter-division time is accompanied by the increase of the added size on a faster time scale than the growth rate. Note however that in our shifts the dynamics of ppGpp levels may follow a two-time-scale dynamics [46, 47].

### A threshold accumulation model of a signal produced at the same rate as a protein expressed from a terminus-proximate constitutive promoter explains the shift division dynamics

We figured that, instead, one may need a model where division dynamics is coupled to the mechanistic action of a biological circuit able to sense the physiological state of the cell. We therefore sought to define a mechanistic model where a protein under physiological control could act as the trigger defining cell division. In particular, we focused on a class of “threshold accumulator” models that have been proposed several times in the literature, both for replication and for cell division [1, 9, 23, 24, 32]. We supposed that the fast and complex changes in division rate and added size observed during our shifts could be due to a coupling between the changes in biosynthetic “sectors” [48] occurring during the shift and cell-division dynamics. The literature offers models that describe proteome sector dynamics and biosynthesis in non-steady regimes [33] and recent attempts were put forward to link these sectors with cell division [49–52]. However, the descriptions differ, and current data do not allow to select a specific one. Thus, rather than committing to a specific choice, we decided to take an experimentally driven approach to define our model.

Following the literature, we agnostically assumed that an effector molecule that is produced at a rate proportional to volume triggers division once a critical amount is reached (the above-mentioned “threshold accumulation” hypothesis). Steady-state data from our experiments provide validation of the fact that protein production is proportional to cell volume (SI Fig. S3), and thus that a volume-specific rate *r* is a good descriptor of protein synthesis from our promoters [53]. Finally, a well-known sufficient requirement to obtain near-adder correlations from an accumulator model is that the trigger molecule is reset to zero upon birth [1, 31, 32]. This last point can be achieved biologically if the divisor molecule is structural, e.g. cell surface material that is integrated into the cell cap upon division [31].

Assuming a putative divisor protein is produced with rate *rV*, proportional to volume (here *r* has units of inverse volume and time), *dN/dt* = *rV*. There is a critical amount *N ^∗^* that once reached, triggers division, *N* (*τ*) = *N ^∗^*, and at cell birth, the number of molecules are reset to zero *N* (0) = 0. Solving for *N*, under steady growth, one obtains

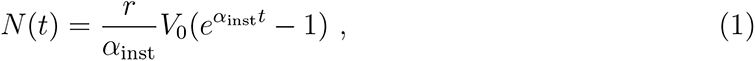

and, as a consequence

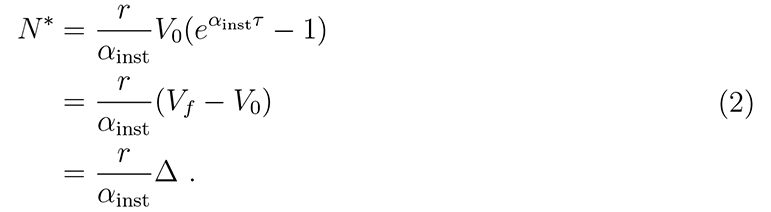

Our strains carried fluorescent reporters for ribosomal and constitutive promoters inserted in the chromosome close to either the origin or the terminus of replication (see methods and ref. [54, 55]), and we used their experimentally measured dynamics to define the putative production rate of the putative “adder protein”. The constitutive reporter “P5” is based on a strong exogenous promoter with consensus −10 and −35 sequences lacking any transcription factor regulation, as well as lacking a GC-rich ppGpp discriminator region at the transcirption initiation site that would render it sensitive to inhibition by ppGpp. Conversely, the ribosomal reporters “P1”, is derived from the P1 promoter of the ribosomal RNA operon rrnB and therefore contains a GC-rich disciminator region at the transcirption start site. The upstream end of the promoter has been omitted, thus deleting the binding sites for any transcriptional regulators (see methods). Thus, we define “constitutive” here as lacking transcription factor regulation as well as lacking a ppGpp discriminator region. In order to to determine the role of ppGpp in the regulation of gene expression and of a possible trigger factor during a growth shift adaptation, we have compared the production rate of GFP in our strains having chromosomal reporters for ribosomal (P1) and constitutive (P5) promoters, and measured the average volume-specific rate, *r*(*t*), across the shift.

A comparison of Eq. (2) with steady-state data for all promoters indicates that *N ^∗^* was about the same in both conditions, before and after the shift (SI Fig. S4). We therefore reverted for simplicity to a model where *N ^∗^* was constant (the data do not allow to quantify this number in absolute terms due to the arbitrary unit of fluorescence measurement). We then ran simulations using as input from experimental data (hence with no free parameters) both the measured production rate *r*(*t*) (for the four strains with reporters) and the growth rate *α*_inst_(*t*) (Fig. 4A). Fig. 4BCD shows that using similar dynamics to the terminus-proximate constitutive promoter P5ter gives very satisfactory agreement with the data, and captures all the complex changes of both interdivision time and (added and initial) cell-size dynamics. For consistency, we also verified that both model and data comply to an expected constraint involving strength of size control, mean size and growth variables (SI Fig. S5).

**FIG. 4.**
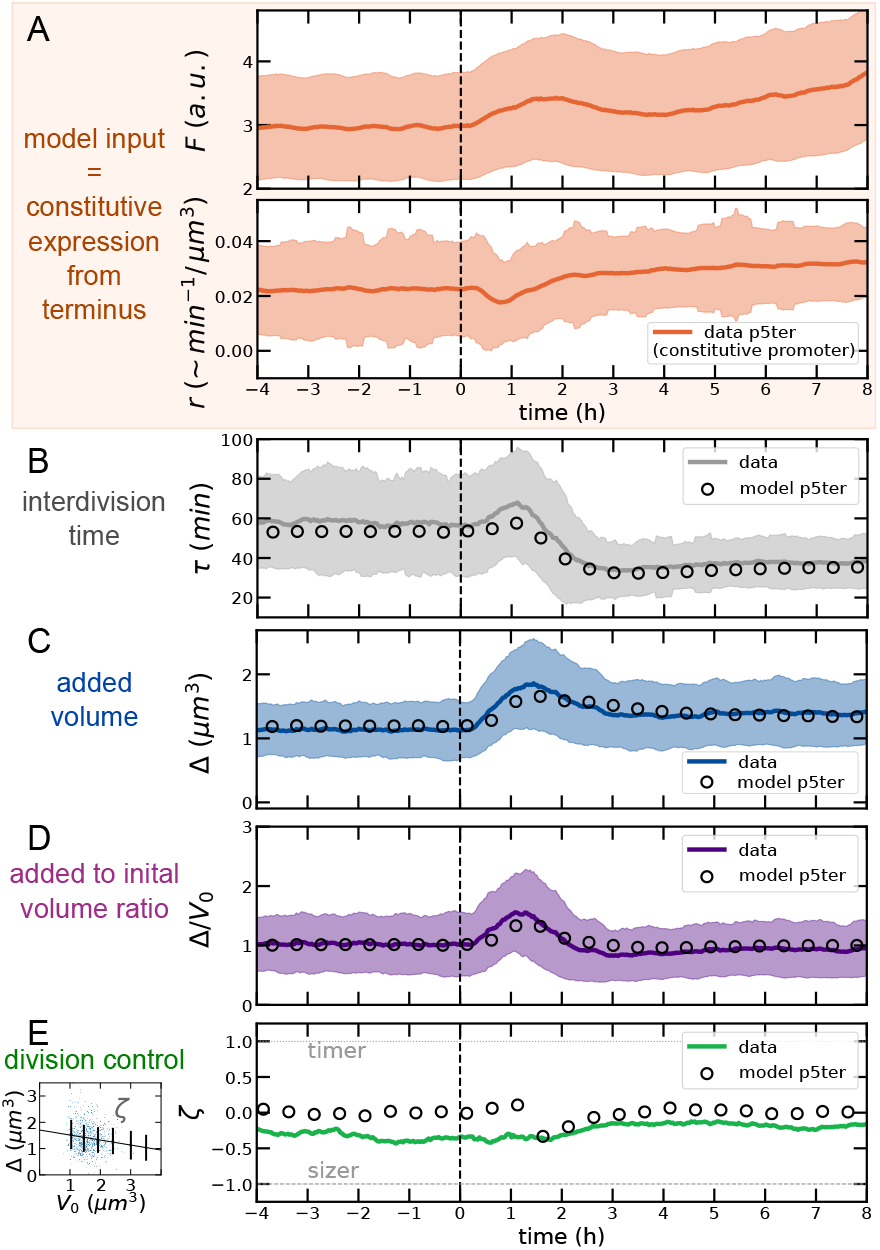
A putative divisor protein expressed from a consititutive promoter explains the shift data. A) As model input we used the measured instantaneous growth rates and volume-specific production rate *r* (obtained from derivatives along lineages) from our promoters, (in this case the P5 constitutive promoter inserted close to the replication terminus). Note that this quantity has units min*^−^*^1^/*µ*m^3^, minus a constant conversion factor from fluorescence to molecule number. The panel also shows the absolute fluorescence *F* from the same promoter. BCD) The model predicts faithfully the size dynamics. E) The model reproduces the observed robustness of near-adder size control. Other model variants using the production rates of different reporters fail to reproduce the observed size dynamics (SI Fig. S6, S7, S8).

While the robustness of the near-adder behaviour across the shift is captured by all the models based on four strains used here (Fig. 4E and SI Fig. S6, S7, S8), the only model that can reproduce the cell size and division time dynamics is the one based on the P5ter promoter. Conversely, the alternative models (based on ribosomal promoter P1 close to both replication origin and terminus, as well as on the same P5 promoter close to the origin) fail to reproduce the size and division time dynamics, reinforcing the idea that the agreement of the data with the model based on the P5ter promoter is not a coincidence (SI Fig. S6, S7, S8). Finally, compared to the models described in Fig. 3 this accumulator model requires only one free parameter (*N ^∗^*, calibrated to match the initial size before the shift). Hence in this case, the steady-state size after the shift is a prediction, which the model performs accurately (validating the idea of a constant critical threshold for the putative adder protein).

### The production rate of the accumulator model can be reverse-engineered from division data, and behaves as a constitutive promoter

An independent reverse argument on the same model leads to the same conclusions, without using the promoter expression data as model input. This argument assumes that *N ^∗^* is constant after the shift, and determines the volume-specific rate that the putative adder molecule needs to have in order to reproduce the data. Solving for *r* in Eq. (2) gives a requirement on the volume-specific transcription rate in steady growth. In the Materials and Methods we generalize Eq. (2) out of balanced exponential growth, when the instantaneous growth rate *α*_inst_(*t*) and the volume-specific rate of expression *r*(*t*) are time-dependent. The full equation cannot be analytically inverted to infer *r*_reverse_(*t*) from the instantaneous growth-rate *α*_inst_(*t*) and the added size Δ(*t*). Nevertheless, the numerical solution matches very well the following expression

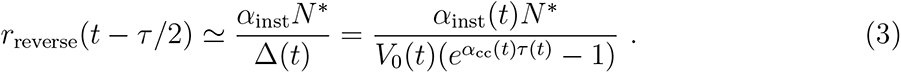

The delay comes from the fact that the integrated production rate over the cell cycle causes the (future) added size at the division time. This effect is fully captured by the expression presented in the Materials and Methods.

SI Fig. S9 shows the behavior of *r*_reverse_ in the transient period after the shift, which initially decreases and then reaches a higher plateau than found before the shift. This behavior closely resembles the dynamics of the P5 constitutive promoter. The precise value of *N ^∗^* does not affect this “down-up” dynamics, and just sets a quantitative vertical offset in the plot shown in SI Fig. S9; it is important for this reverse argument however that *N ^∗^* remains constant after the shift. Comparing SI Fig. S9 with Fig 4A confirms that constitutive expression meets the requirements set by our shift data for the accumulated divisor factor. We also note that the model clearly works well in a fit where the threshold is different before and after shift, and in this case the best-fit parameter values are such that *N ^∗^*(pre-shift) *,.. N ∗*(post-shift), which confirms that the value of the threshold remains constant or nearly constant.

The model obtained by the reverse argument shows an even more faithful agreement with data (SI Fig. S10). We also applied this model to an upshift experiment between the same two nutrient conditions at 30*°*C (SI Fig. S11), finding that the inferred rate by the reverse model shows the same pattern over the shift, initially decreasing then increasing with an overshoot, before reaching a plateau. Finally, we used the reverse model for more detailed comparisons with data. First, we analyzed the shift dynamics by a discrete time variable representing the number of generations separating each lineage from the shift. The over-shoots visible in Fig. 2 persist when the same data is plotted by aligning the cell generations before and after nutrient shift, and are captured by both the positive and the reverse model (SI Fig. S12). Second, we performed a detailed analysis of the generation seeing the shift and the subsequent one (SI Fig. S13-S15). This analysis shows the the division behavior of the generation seeing the shift depends on the cell-cycle time at the time the nutrient level changes. The model reproduces well the cell-cycle behavior of the generation seeing the shift and the subsequent ones in terms of interdivision time and added size. However, a careful analysis also reveals discrepancies that are likely due to ingredients not present in our model (and not monitored in our data), such as the interplay of the shift time and the processing of the chromosome during the cell cycle [2, 7]. Specifically, in the model, cells that see the nutrient shift late in their cell cycle do not modify their cycle duration compared to the pre-shift conditions, while in the data these cells delay their division (Fig. S15)

## DISCUSSION

This work has two main components.

First, it examines a well-defined upshift where the carbon source remains the same, and the cell adapts to the presence of the added amino acids, in real time and in single cells, extending classic results obtained when single-cell tracking was not possible [38]. The response to the shift is unexpectedly complex, challenging old conclusions from population analysis. Most variables, including growth rate, show a two- or three-phase dynamics, with trend changes, overshoots and undershoots before they fully relax; growth rate is slow to achieve full relaxation to its nutrient-imposed value; interdivision time and added size start changing early on, in parallel to growth rate itself (and not slaved to this variable). A near-adder correlation is maintained throughout the shift despite the complex response. We are aware that much of this complexity will need deeper theoretical and experimental investigations to be fully unraveled, but our data set could set an important reference for forthcoming studies. Here, we decided to restrict our focus on the division-control dynamics, and validation of competing cell-division models.

Second, the unexpected kinetics during the shift are qualitatively incompatible with several current models for control of cell size in single cells. This holds irrespective of the details. Thus, a new way of looking at the cell division decision upon a shift in growth conditions is required. We explore experimentally the possibility that division could be triggered by the accumulation of a critical amount of a constitutively expressed (i.e. presumptively not growth-rate regulated, or not directly) molecule. As we show, the production levels for such a molecule explain the unexpected division kinetics is if the molecule is expressed from an unregulated promoter located near the terminus.

Near-adder size-control behavior is robust across the shift, despite of large changes in the cell-cycle parameters as well as in the correlations between initial size and growth rate in this transient regime. This feature is surprising on biological grounds, and as we show, it is a common feature of the current phenomenological models, where it is built in the assumption that either the inter-initiation or the inter-division process invariably should reproduce adder correlations (for the case of an initiation sizer, we observe the same constant behavior, with a different value of the size-control parameter). This common assumption is essentially post-hoc when looking at steady-state data, but its applicability to transient data indicates that its validity might be broader. Hence, we take the robustness of the size-control parameter across the shift as a validation (beyond expectation) of the widespread approach to modeling size control.

However, not all models can reproduce the complex temporal changes observed in the key cell-cycle parameters monitored here, and specifically of interdivision time, initial size, and added size. Standard frameworks describing cell size control [13, 14], are “hierarchical”, in the sense that (constant, or fluctuating) nutrients are thought to affect growth rate, and size and division rate in turn adjust to the growth rate [45]. However, division rate may sense nutrients directly [45], and become decoupled from the different cell growth processes connected to the nutrient-imposed growth rate (bulk mass biosynthesis, elongation, volume change). It is worth noting that, as soon as the nutrient condition is changed, both added size and interdivision time show changes before growth rate begins to change and reach a new steady state before growth rate does, thus invalidating a view where the cell cycle “senses” growth rate and adapts the interdivision time accordingly. A scenario where both the division and the biosynthesis machinery are able to sense and react to nutrient changes appears more plausible. As we further show, the standard view where cell division is slaved to the chromosome replication-segregation cycle appears to be at odds with the observed time scales and qualitative changes in cell-cycle parameters across the shift. Both results are in accordance with a parallel study on downshifts induced by increased ppGpp [45]. Beyond the tested models, the same limitations may be common to most models available in the literature, (which were not designed to describe shifts) as a consequence of the hidden assumption whereby these models describe division rate as a consequence of growth rate, treated as an input parameters [5].

Finally, using the data from four GFP reporter strains with either a ribosomal or a constitutive promoter inserted at two different locations along the genome, we could test the hypothesis that the adder molecule is a protein under the control of these kinds of promoters (whose expression is differently dependent on growth rate). We believe that, giving our current limited knowledge on the crosstalks between growth- and cell-cycle physiology, this “empirical” approach to forward models is preferrable to more theoretical routes postulating the behavior of different proteome sectors across shifts [49–52]. Comparing the predictions of the four possible models that can be generated from our data, we show that the data are in agreement with a model where a putative adder protein is under the control of a constitutive promoter located close to the replication terminus of the *E. coli* chromosome. This model predicts efficiently the behavior of interdivision time, initial size and added size of cells across the shift. Conversely, if we reverse-engineer the production rate of an accumulator model in order to reproduce our division data, we find that it matches the expression pattern of the ter-proximate constitutive promoter.

Crucially, the specific production rate of a constitutive promoter transiently decreases immediately after the upshift, while the one of a ribosomal promoter transiently increases, in agreement with the rapid decrease in ppGpp upon addition of amino acids to cells grown in their absence [36]. Note however that in this study we do not investigate *why* the observed promoters show this complex behavior, which is a separate question [33]. Presumably, cells are spending their resources producing more ribosomes to take advantage of the richer media to such a level that constitutive promoters are fractionally downregulated. We also note that surface and mass synthesis decouple during nutrient shifts [56].

The conclusion that a threshold accumulation process sets division is common in the recent literature (to the point that isolating competing hypotheses is an open challenge). A recent study combining experiments and theory concludes that in order to reproduce the observed trends across conditions, this molecule should be under growth-rate control and able to sense the mean duration of the replication or segregation period of the chromosome in a given condition [10]. Several theoretical studes have speculated on the possible interactions between biosynthetic “sectors” and cell-division control [49–52]. In particular, based on the analysis of steady-state growth-division data, two studies [49, 50] have suggested that the molecule effecting the adder behavior may be a protein of the “P sector” of the proteome, i.e., a constitutive promoter, in full agreement with the conclusions reached here.

It has been speculated that the divisor molecule could be FtsZ itself [9, 32]. In agreement with our findings, FtsZ expression is not ppGpp-dependent [57]. Additionally, its expression rate is inversely proportional to the replication/segregation period duration [9, 10, 49]. However, the subcellular localization dynamics and activity of this protein are complex [58]. Additionally, the dynamics of width across the shift in our data (SI Fig. S1) is very slow compared to the changes in division rate and added size. Hence, our data seem at odds with a scenario whereby the FtsZ ring (or the septum) are always rate-limiting for cell division, which predicts a direct link between cell diameter and division rate [31, 32].

A parallel single-cell study has implicated the alarmone ppGpp [45], the key player of a network sensing nutrient levels and setting the growth rate through transcriptional control of ribosomal promoters, in the *direct* regulation of division rate. This result is based on the observation that the added size and interdivison time change faster than the growth rate in a downshift induced by upregulation of ppGpp synthesis. This observation opens the question of whether ppGpp could have direct or indirect interactions with the divisome.

In our shifts, the carbon source does not change, and available ppGpp levels are expected to decrease due to the increased levels of amino acids in the new media [46], so the nutrient change could be paralleled to an effective rapid decrease in ppGpp levels, similarly to the case studied by Buke and coworkers (in our case, we also expect that the transcription of genes for AA biosynthesis is shut off, freeing resources for the increased expression of genes involved in the transport of AA and biosynthesis of cellular components). Based on our data and modeling analysis, we speculate that the control of ppGpp on cell division could be exerted through an indirect effect of the change in ppGpp on the transcription of a constitutive promoter, supporting the idea that the more direct ppGpp-divisome interaction proposed by Buke and coworkers might not be necessary.

Our data and model show that this transcriptional effect can explain the fast time-scale changes observed in the shift. ppGpp binds directly to RNA polymerase (RNAP) and destabilizes its interaction with the promoter, with a strong effect on the transcription initiation rate of ribosomal promoters, where the RNAP-promoter complex is particularly weak [46, 59]. A rapid decrease in ppGpp could thus result in an increase in the amount of RNAP that is used to transcribe ribosomal genes, decreasing the amount of RNAP that is available for the transcription of other promoters, such as the P5 promoter used here. Hence, a change in ppGpp levels has in principle the power of changing the production rate of most gene categories on relatively fast time scales [36, 48]. Our analysis shows that this action could be sufficient to modulate division rate very quickly across an upshift, and faster than effecting the observed changes in overall growth rate. As a rationale for this, we hypothesize that a change in the overall biosynthetic rate requires production of many ribosomes, whereas a sufficiently low threshold of a putative “adder protein” could make the effects of a change in its production rate on the cell division rate very quick to observe.

To conclude, we propose a more generic and formal interpretation of our main results, which may help guide future analyses aiming to identify molecular players of cell division. The accumulator model can reproduce the data if two crucial prescriptions are included. First, the production rate has to transiently decrease during the upshift, then increase, in order to capture the behavior of the division rate. Hence, the divisor molecule must be something that responds in a precise way to changes in ppGpp levels, without using biosynthesis as a readout. Second, since the terminus-proximate location of the promoter appears to play a role, any putative molecular accumulator circuit leading to cell division should be able to count terminus copies or something correlated.

## MATERIALS AND METHODS

### a. Experimental details Strains and culture conditions

The six strains assayed here each contain one of three promoters controlling GFP expression from the gfpmut2 gene [60] at either a genomic position close to the origin or the terminus of replication, with *E. coli* K-12 BW25113 serving as the background. The former is at genome coordinate 4413507 bp, between the two convergent genes aidB and yjfN, while the latter is at 1395706 bp, between the two convergent genes ynaJ and uspE. The two promoters used here correspond to the following: the first contains a shortened version of the well-characterized ribosomal RNA operon promoter rrnBP1 [61, 62], here called P1, which includes the sequence from −69 to +6 relative to the transcription start site. The binding sites for Fis and the higher affinity H-NS binding site are thus omitted from this construct. The rrnBP1 promoter has a GC-rich discriminator region at the transcription initiation site, which makes the open complex sensitive to changes in negative supercoiling and to inhibition by ppGpp [62]. The second promoter used here is a constitutive promoter, P5, derived from the phage T5, that has consensus −10 and −35 sequences and no discriminator region. The strong constitutive P5 promoter comes from the bacteriophage T5, while the growth-rate dependent ribosomal promoter rrnB P1 normally acts on the *E. coli* rRNA operon. Per strain, one of the above promoters was placed at one of two chromosomal locations: either close to the origin of replication at ori-3 (4413507 bp, nearest gene aidB); or the replication terminus at ter-3 (1395706 bp, nearest gene ydaA). A kanamycin resistance cassette (KanR) is divergently expressed from each promoter-gfpmut2 cassette. Bacteria were always kept at 37*°*C. Growth media had M9 minimal medium (BD Difco, Wokingham, UK) and 0.4% glucose glucose as the carbon source; the upshift consisted in switching M9-glucose with M9-glucose supplemented with 0.5% casamino acids (Sigma-Aldrich Company Ltd., Dorset, UK; OmniPur). Overnight cultures were diluted 500:1 in new, slow-growth medium and returned to the incubator for 3-4 h, i.e. until the cells reached their exponential phase. 1mL of culture was then pipetted into two 2mL prewarmed Eppendorf tubes and centrifuged at 4000 rpm and 37*°*C for 4 min. The supernatant was pipetted off and each pellet was resuspended in 40-50 *µ*L of fresh slow growth medium by vortexing. These condensed cultures were then injected into prewarmed, passivated and rinsed PDMS chips as described below.

### Device fabrication and loading

PDMS-based (Sylgard by Dow Corning, USA) microfluidic devices were constructed to support the long-term growth of *E. coli* under a nutrient upshift from slow to fast growth medium, and bonded onto glass coverslips (Menzel-Gläser, Germany; 22 x 60 mm), as described in [35]. PDMS was cast over the master template to yield the negative relief mother machine pattern to be fixed to the coverslip. Before loading bacteria into the device, each chip was treated with a solution of bovine serum albumin (BSA; Sigma-Alrich Company Ltd., Dorset, UK) to minimize bacterial interactions and binding to the glass or PDMS components. Devices were pipetted with 5 *µ*L of 10% BSA and incubated at 37*°*C for 45-60 min. Passivated chips were rinsed with fresh medium. Bacteria in exponential phase were then similarly pipetted into the device. The chip was then mounted on a spin coater (Electronic Micro Systems, Sutton Coldfield, UK; Model 4000) such that the line connecting both injection sites intersected the axis of rotation. Each sample was spun at 3250 rpm for 10 minutes and bacteria were pulled into the microchannels by the resulting centripetal force. Chips were inspected under 40x magnification to confirm that a substantial fraction of microchannels contained bacteria. In the case of poor trapping, the sample was spun again under the same settings.

### Microfluidic circuit and growth medium switch

Two 20 mL syringes (Becton, Dickinson UK Ltd., Wokingham, UK; Plastipak) were prepared with each growth medium and mounted on separate syringe pumps (KD Scientific, Holliston, USA; Legato 110). A membrane filter (Sigma-Aldrich Company Ltd., Dorset, UK; MF-Millipore, 0.22 *µ*m pore size) was placed between the syringe and the dispensing tip (Intertronics, Kidlington, UK; stainless steel, straight blunt, 1/2”, 23 gauge), ensuring that only sterile growth media entered the microfluidic device. A Y-junction was formed with three pieces of Tygon tubing (Cole Parmer, St. Neots, UK; 0.020” x 0.060” OD; 40, 40, and 20 cm lengths) and a PEEK Y-connector (Kinesis, St. Neots, UK; for 1/16” OD tubing, flangeless fittings). The two 40 cm tubing segments were attached to the syringe dispensing tips and the 20 cm tubing segment was connected to the input injection site of the chip using the stainless steel shaft of a 90 degree dispensing tip (Intertronics, Kidlington, UK; stainless steel, 90 degree bend blunt, 22 gauge). Prior to attaching the 90 degree connector to the chip, the fast-growth medium was pumped through the tubing at 100 *µ*L/min until about 0.5 mL had drained from the connector end. The same was then repeated using the slow growth medium. The syringe pump controlling the slow growth medium was then set to 10 *µ*L/min and the input connector tip was inserted into the chip. The two syringe pumps were programmed to alternate so that the slow-growth medium fed the device first, while the fast-growth medium pump was delayed. In most cases, a complete upshift experiment lasted between 18 and 24 h with roughly equal time spent in each growth medium. While the chip was plugged in, the infusion flow rate was set to 10 *µ*L/min irrespective of growth medium. This corresponds to a flow velocity in the main feeding channel of approximately 166.7 mm/s. There was a brief delay of roughly 4 min before the new medium reached the device, which was accounted in our definition of the shift time. This corresponds to the 20 cm distance that the fluid must travel from the Y-connector junction to the chip. We also performed an experimental test of the time to fill a channel and the device itself, by flowing (fluorescent) LB media (SI Fig. S16). It takes less than 5 minutes (one frame) to completely fill a channel with medium. The mixing of the two media outside the channels is negligible judging from the fluorescence, as expected from such a low Reynolds number flow. Additionally, we tested directly in the data that the growth rate averaged on the channels at the two outermost sides of the device shows a small delay, of about 5 minutes (SI Fig. S16).

### Optical microscopy and image acquisition

Images were acquired under 60x magnification using an inverted microscope (Nikon Eclipse Ti-E, Tokyo, Japan; oil objective, NA 1.45). The camera (Andor iXon DU-897 EMCCD, Oxford Instruments Industrial Products Ltd, Abingdon, UK) captured 16 bit images at 512 x 512 pixel resolution with the length of one pixel equal to 0.1067 *µ*m. The motorized stage and camera were programmed to cycle between at most 40 fields of view, each spanning roughly 8 microchannels, every 5 min, recording three images per field of view: one brightfield (red or green illumination; excitation PD01 and PM01, LUXEON Z LED, Lumileds, Schipol, Netherlands; multiband FF01-392/474/554/635 optical filter, Semrock, Rochester NY, USA), one fluorescence (blue, PB01), and one blank without illumination. The exposure time for all images was 0.286 sec, corresponding to the cameras maximum shutter speed. All LEDs were set at their respective minimum forward currents to minimize potential interference with the cells due to overexposure. The entire apparatus was left to acquire images continuously through the growth medium switch, with sufficient data before and after the switch event. The coverslip of the microfluidic device was fixed onto a custom aluminum thermoelectric cooler (TEC) that was then fitted on the microscope stage. The stage and objective temperatures were always set to 37*°*C. The input tubing was woven through inlet holes to ensure that fluid entering the chip was the appropriate temperature.

### b. Data analysis details. Segmentation and tracking algorithms

We developed bug-pipe, a custom-built MATLAB package to process mother machine-based images and handle the resulting cell data. In brief, for each background-subtracted fluorescence image the segmentation algorithm proceeds in the following steps: 1) find regions of interest (ROI), i.e. the channels trapping each cell line; 2) within each ROI, threshold and isolate cells in the background; 3) find cell boundaries and further process any “fused” cells; 4) remove artefacts based on size. Simple background subtraction is performed by subtracting the respective dark frame from each fluorescence image. The tracking procedure that links individual cells between consecutive frames takes advantage of the geometric constraints imposed by the microchannel format of the microfluidic chip. Namely, that in the absence of cell divisions, a cells rank within a channel is conserved between frames. The tracking procedure operates by: 1) microchannel matching between frames, 2) sorting cells in-channel based on distance from the dead-end, and 3) determining markers for cell division to adjust the rank-based pairing as necessary. The package was designed with separate functions for segmentation, tracking, and data handling procedures. This grants flexibility for the user if, say, only big data manipulation tools are needed. Detailed information on bugpipe is available as SI Information. The bugpipe package and its complete documentation is available at https://github.com/panlilio/bugpipe. The data analyzed here refer to 3 repeats of the entire upshift experiment for the P5ori promoter, 2 repeats for the P5ter and P1ori promoters and 1 repeat for the P1ter promoter.

### c Modeling details

We considered four alternative models to describe cell-size control in steady growth and during the nutrient shift. These models (with one exception) are known to reproduce the adder correlations in steady growth and can be parameterized to reproduce the empirical relation between size and growth rate across condition, known as Schaechter’s law [10, 37].

We generalize these models to non-steady conditions under nutrient shifts. This generalization requires non-straightforward decisions. Here we employ a data-driven strategy and use the experimental trajectory of the average growth rate *ᾱ*(*t*), and of other quantities if applicable, as input of the model. The model output that we test against the experiments are the trajectories of division time, added size, initial size, and strength of division control during the shift.

In order to take into account for the non-steady values of the single-cell growth rate *α*_inst_, we consider the following dynamics

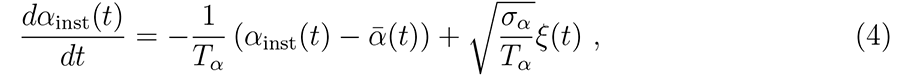

where *ξ*(*t*) is a Gaussian white noise with zero mean and a standard deviation equal to one. Under constant values of *ᾱ*(*t*) (*ᾱ*(*t*) = *ᾱ*), this equation corresponds to a Gaussian stationary distribution of the *α*_inst_(*t*) with mean *ᾱ* and standard deviation *σ_α_*. In this case, the parameter *T_α_* sets the autocorrelation time of the single-cell growth rates. When instead *ᾱ*(*t*) varies, *T_α_* sets the typical time that it takes for a single cell to respond to the forcing imposed by *ᾱ*(*t*). It is easy to show that the average growth rate is equal to

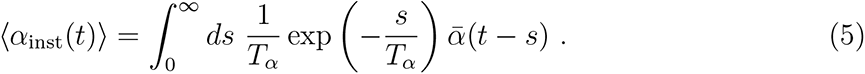

In order to simplify the inference of *ᾱ*(*t*) we assumed *T_α_* to be small compared to a cell cycle and to the typical time of change of *ᾱ*(*t*) during the nutrient shift. Under this assumption we set *ᾱ*(*t*) = 〈*α*_inst_(*t*)〉. For numerical purposes we set *T_α_* = 1 min. We also verified that our simulations gave the same results extracting the values of *α*_inst_ from the experimental values in each time bin.

### Initiation-sizer model

The model assumes that initiation starts at a constant cell size per origin of replication (“initiation mass”), i.e. *(V_B_)* = *OV ^∗^*, where *V_B_* is the size of the cell at initiation, *O* is the number of origins, while *V ^∗^* is a constant size (independent of the condition) [43]. Division happens at a time *(t_C_*_+*D*_〉 after initiation. In steady growth the model reproduces Schaechter’s law and a sizer-like division control. This model is different (simpler) from the one used in ref. [30], which includes nontrivial correlations (see [7, 8] for a thorough discussion). Single-cell growth rate trajectories were modeled as explained above. We set the two other free parameters of the model (*V ^∗^* and *(t_C_*_+*D*_〉) to reproduce the initial average size at the two steady growth conditions before and after the shift. In particular, in steady growth, it is known [10, 30] that *(V*_0_〉 = *V ^∗^* exp *ᾱ(t_C_*_+_*_D_)*, which can be easily inverted to determine *V ^∗^* and *(t_C_*_+_*_D_)* using the values of *(V*_0_〉 and *ᾱ* before and after the shift. Both the initiation and time-to-division are stochastic, where *(V_B_)* and *(t_C_*_+*D*_〉 are averages. We assume Gaussian noise on log *V_B_* and *t_C_*_+*D*_ with equal magnitude, set to reproduce the coefficient of variation of the initial size (which is independent of the condition [21, 39]).

### Incremental model [16]

The model assumes that cells add on average a constant size per origin Δ*^∗^* = (Δ*_I_)/O* between initiations of replication. As in the initiation-sizer model, division takes place at a size-independent time *(t_C_*_+*D*_〉 after initiation. Single-cell growth rate trajectories were modeled as explained above. The two free parameters of the model (Δ*^∗^* and *(t_C_*_+*D*_〉) are set to reproduce the initial average size at the two steady growth conditions before and after the shift using the relationship *(V*_0_〉 = Δ*^∗^* exp *ᾱ(t_C_*_+_*_D_)*. Both the initiation and time-to-division are stochastic, where the log Δ*_I_* and *t_C_*_+*B*_ are independent Gaussian random variable with equal standard deviation, set to match variability of initial size.

### Relative-rates model

This model assumes that septum-formation is the rate-limiting process for cell division [31]. Synthesis of the target surface material *S* happens at a rate proportional to the volume *dS/dt* = *βV* and *S* is reset to zero at every cell division. *β*(*t*) along the shift is obtained from measured cell surface (from width and length, assuming apherocylinder) in our data. Division is triggered when *S*(*t*) reaches a critical threshold *S^∗^*. Depending on the assumption [31], the threshold could be constant or proportional to the septum area *πw*^2^, with *w* being the width of a cell. We chose the former. Since the two models differ in the threshold *S^∗^*, while *beta*(*t*) is the same, they cannot both reproduce the average added size post shift. The former version (constant *S^∗^*) correctly predicts the average added size in steady state after the shit, while the latter does not. Neither version can reproduce the overshoots in our data. The rates *beta*(*t*) before, during, and after the shift is determined from the empirical value of the width *w*(*t*). The rates *r* before and after the shift are the two free parameters and are determined to reproduce the empirical values of the size before and after the shift. Noise is introduced in the value of *S^∗^*(*t*) which has Lognormal fluctuations set to match the variability of initial size.

### Constitutive/ribosomal divisor accumulator model

This model assumes that the accumulation of some protein triggers division. Synthesis of the divisor protein *N* happens at a rate proportional to the volume *dN/dt* = *rV* and *N* is reset to zero at every cell division. Division is triggered when *N* reaches a critical threshold *N ^∗^*. We assume that *r*(*t*) is proportional to the experimental volume-specific rate of expression of a constitutive or ribosomal promoter from our data, which we use as an empirical input of the model, together with the growth rate (both quantities are evaluated as. The only free parameter of the model is the threshold *N ^∗^* which we assume to be constant during the shift and condition independent. We set *N ^∗^* to match the average size at birth before the shift. Note that, contrarily to the other models described above, we do not set any free-parameter to reproduce the initial size after the shift, which is therefore another prediction of the model. Stochasticity is introduced in the value of *N ^∗^*, which is assumed to have Lognormal fluctuations, with coefficient of variation set to match the variability of initial size.

### Reverse variant of the divisor accumulator model

This model is identical to the previous one, but instead of using promoter expression data to define *r*, it uses the production rate inferred from data on added size and growth rate assuming the model. Equation 2, which holds in balanced exponential growth, can be generalized during the shift, under the assumption that the time-dependency of the added size emerges as a consequence of the time dependency of the instantaneous growth rate *α*_inst_(*t*) and of the volume-specific expression rate *r*(*t*), while the threshold *N ^∗^* remains constant.

An average cell born at time *t* with initial volume *V*_0_(*t*), at age *a* will have volume 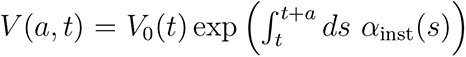. The amount of divisor protein of a cell born at time *t*, with age *a* will be therefore 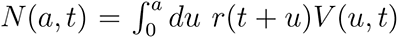. At division *N* (*a_d_, t*) = *N ^∗^*, independently of *t*. From which we obtain

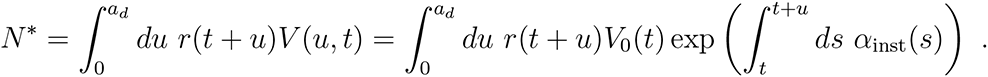

This expression can be written in terms of the average added size Δ(*t*) of a cell born at time *t*, by using the fact that

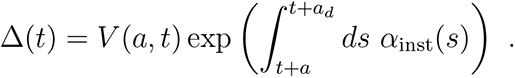

By inverting this expression we obtain the approximate expression for *r*_reverse_(*t*) shown in the main text and in SI Fig. S9. The inferred production rate was compared to the experimental values for the measured promoter ad used in direct simulations of the model.

## ACKNOWLEDGMENTS

This work was supported by the International Human Frontier Science Program Organization, grant HFSP RGY0070/2014. MCL was funded by the Italian Association for Cancer Research AIRC-IG (REF: 23258). We are grateful to Lexuan Liu for help with the experiments, and to G. Micali, F. Büke, S. van Teeffelen, A. Colin, N. Kleckner for feedback on this work.

## SUPPLEMENTARY INFORMATION FOR PANLILIO *ET AL*

### ACCUMULATION AND RESET OF A CONSTITUTIVE PROTEIN TRIGGERING DIVISION EXPLAINS DYNAMICS OF CELL SIZE CONTROL ACROSS NUTRIENT UPSHIFTS

#### SUPPLEMENTARY FIGURES

**FIG. S1.**
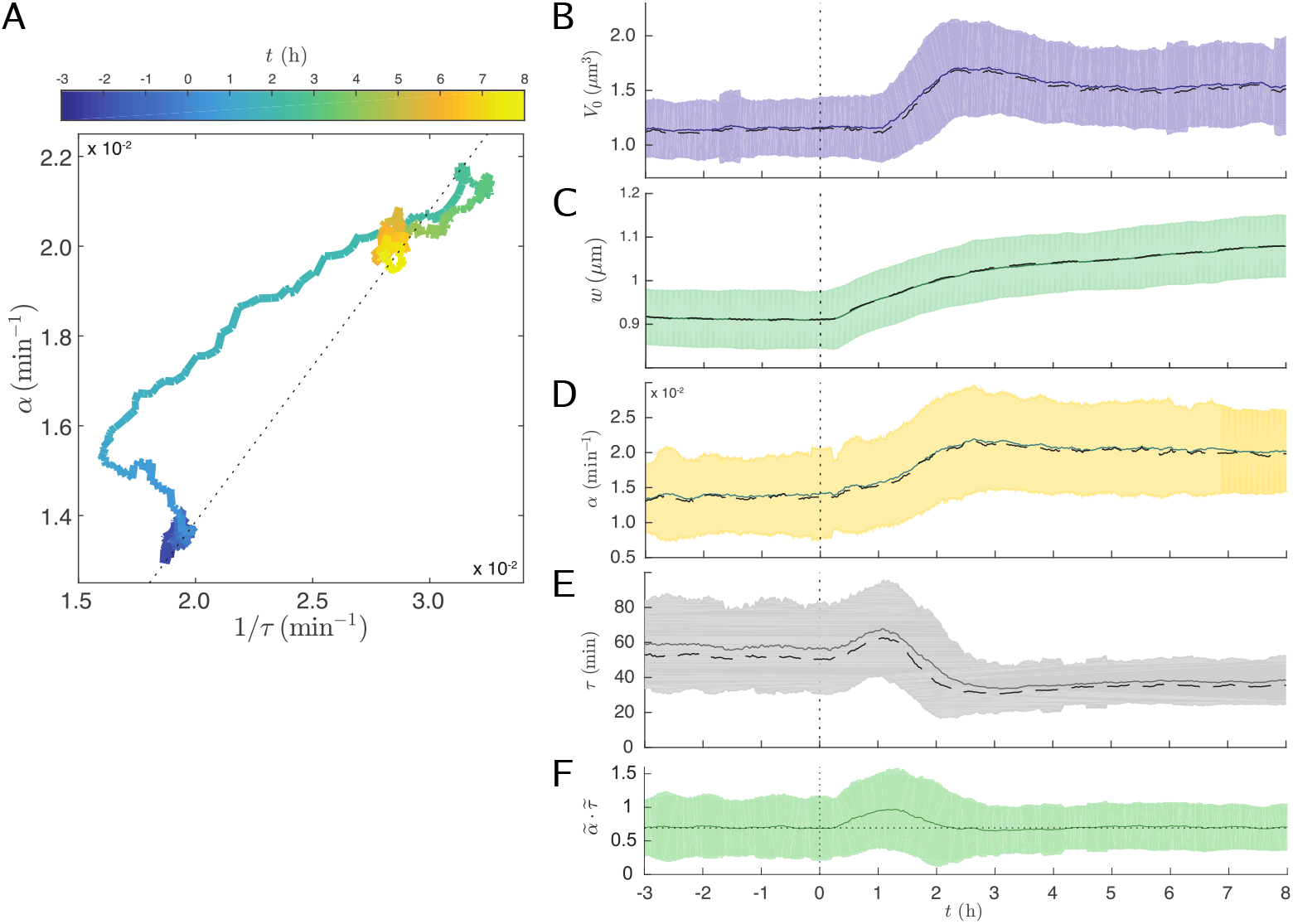
Complex dynamics of growth and division processes across nutrient shifts. A) Time trace in the (mean) 1*/(τ)*-〈*α*_inst_〉 diagram. These two quantities reach fixed points where they are proportional to each other (by a factor of log(2), dotted line) in steady-growth conditions, but they deviate forming a complex pattern during the shift, due to the different time scales involved. B) Mean birth volume initially overshoots its target increased value in rich media. C) Mean cell width relaxes slowly in the new condition. D) Mean growth rate as a function of time into the experiment. E) Mean interdivision time as a functon of time in the experiment. Note that this quantity has an early increasing trend, opposite to the target of the new condition (i.e., cells whose target is to divide faster, initially divide with a slower pace) inreasing trend). F) The product of median growth rate and median interdivision time deviates from a constant value during the shift (“non adiabatic” transition). Media shift time is indicated at vertical dotted line. Unless otherwise stated, solid line indicates observable mean, dashed line indicates median, and shaded regions indicate SD for all time series.

**FIG. S2.**
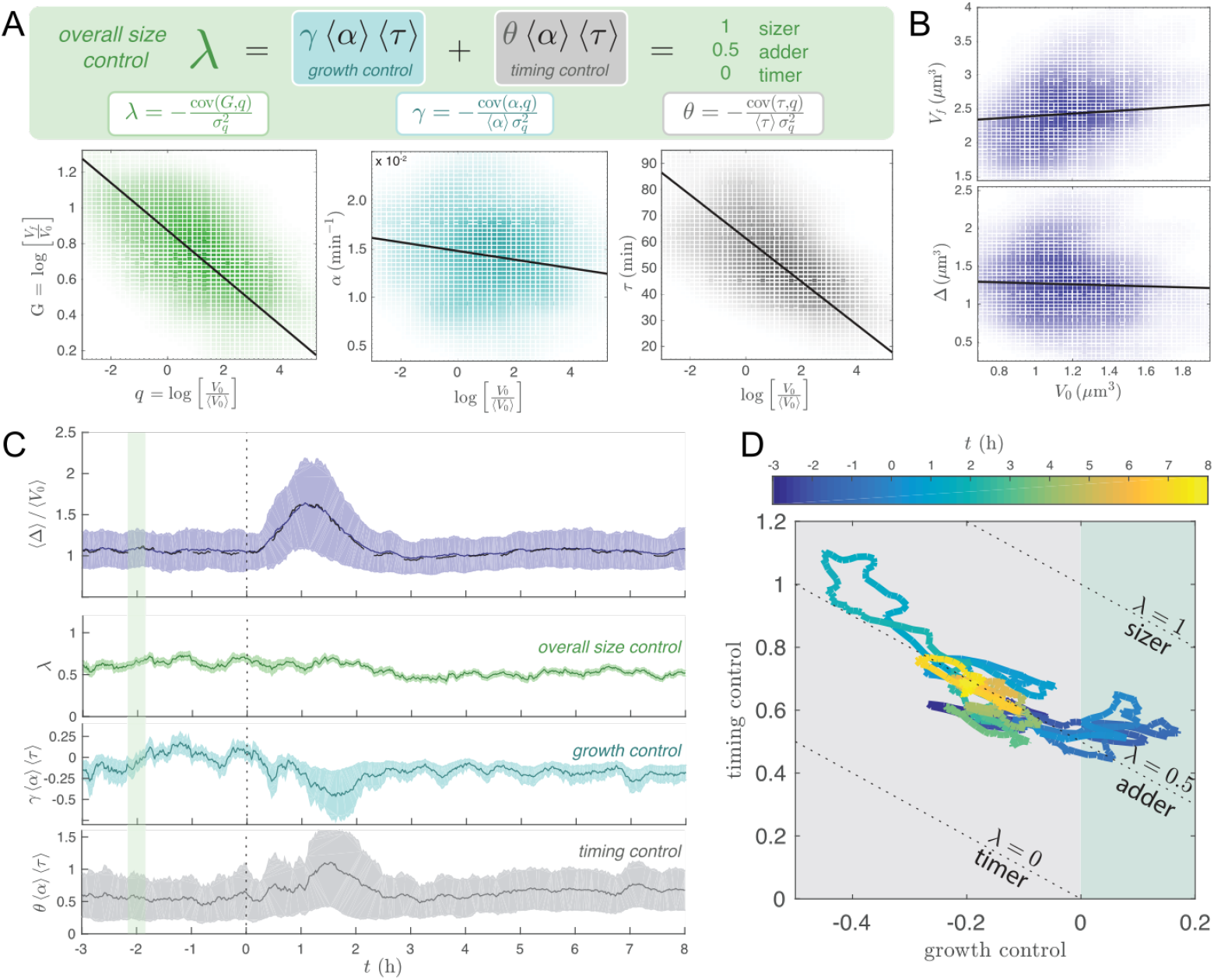
Near-adder behaviour is through shifts is preserved through compensation of division timing control and growth-rate control despite complex dynamics between growth and division processes. Media shift indicated at dotted vertical line. A) The size-growth plot measuring multiplicative growth vs logarithmic initial size quantifies size corrections wuth the slope *λ* (=1 for sizer, 0 for timer, 1/2 for adder) and can be split into inter-division time and growht-rate contributions [17]. B) Equivalent to these slopes, one can measure the slope of the added size or final size *vs* initial size [15, 17]. CD) Dynamics of size correction across the shift. The slope of the size-growth plot *λ* remains constant, and timing/growth control variables compensate to maintain near-adder correlations.

**FIG. S3.**
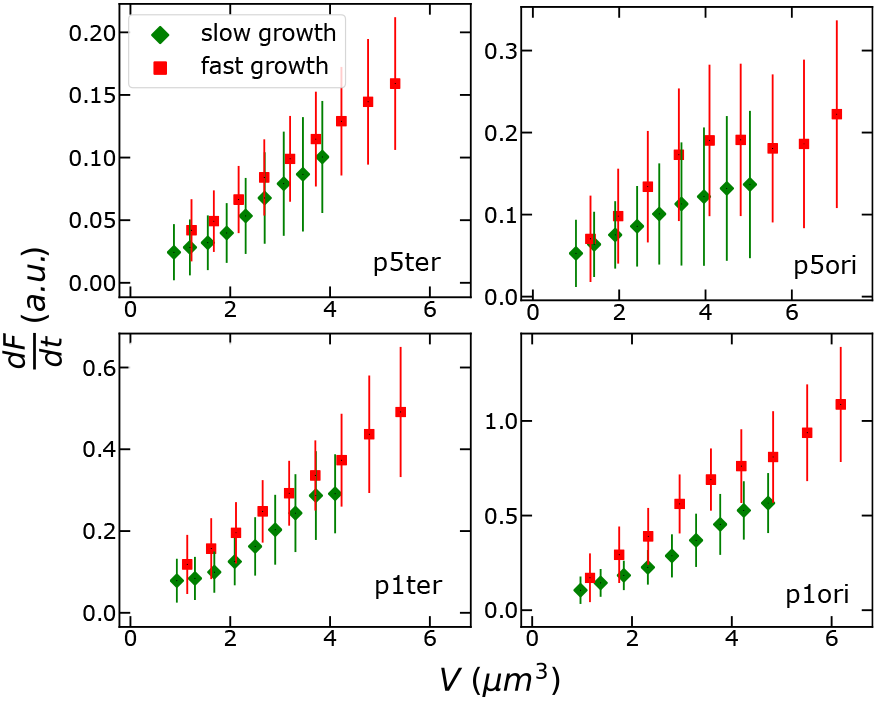
GFP production rate from the ribosomal and constitutive promoters used in this study is proportional to cell volume. The data shown here come from measurements of GFP expression from the P5 and P1 and promoters in different chromosomal locations (different panels) in slow (M9 + 0.4% glucose, green diamonds) and fast (M9 + 0.4% glucose + 0.5% casamino acids, red squares) steady growth conditions. Points on the *y* axis are averages of discrete derivatives of fluorescence-versus-time tracks *dF* (*t*)*/dt*, correlated to measured volume (*x* axis). Error bars indicate the standard deviations on binned averages.

**FIG. S4.**
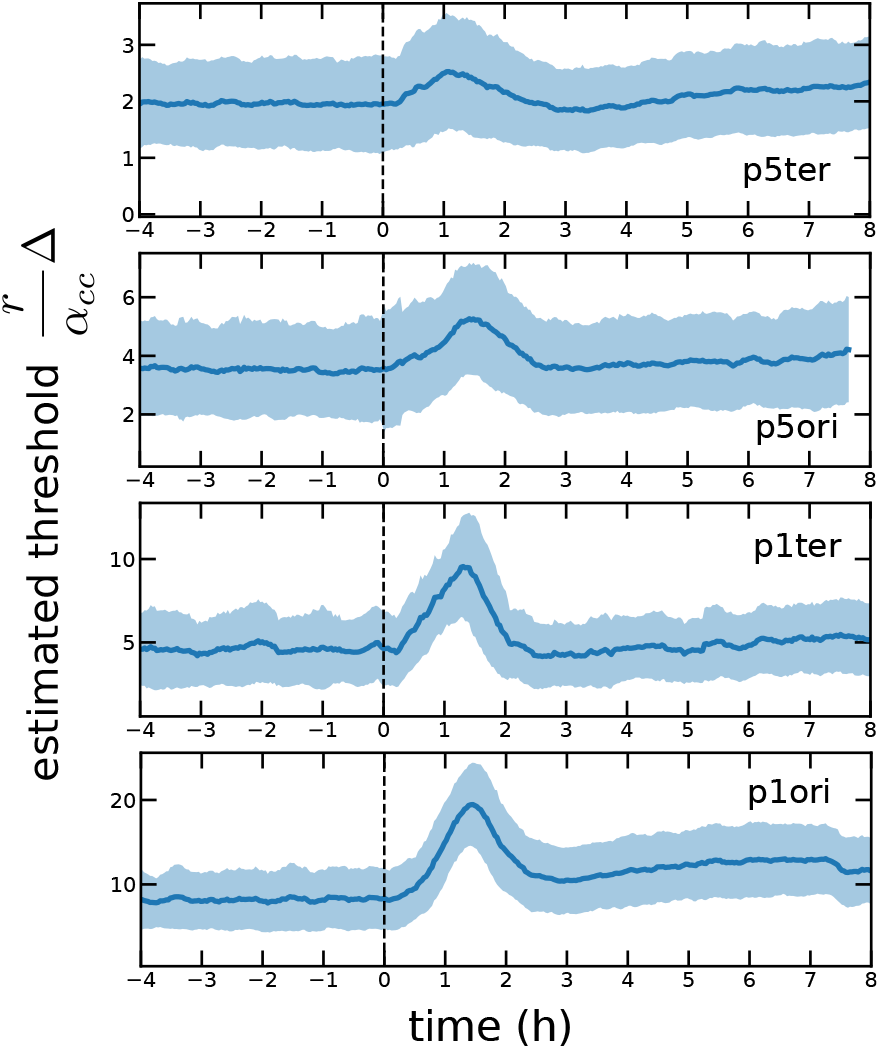
The estimated threshold for an accumulator model is similar across conditions for all tested promoters. Assuming the accumulator model described in the main text, the plot evaluates a running average of the quantity 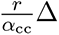, which, at steady state, corresponds to the theoretical prediction of the threshold value *N^∗^* of the accumulator model (see Eq. (2) of the main text). The data are compatible with a threshold that remains roughly constant across the two conditions. The production rate *r* = 1*/V dF/dt* is averaged over cell cycles, and the growth rate *α*_cc_ comes from exponential fits of the volume *vs* time data of the same cell cycle. Both variables are associated to the division time of a cell.

**FIG. S5.**
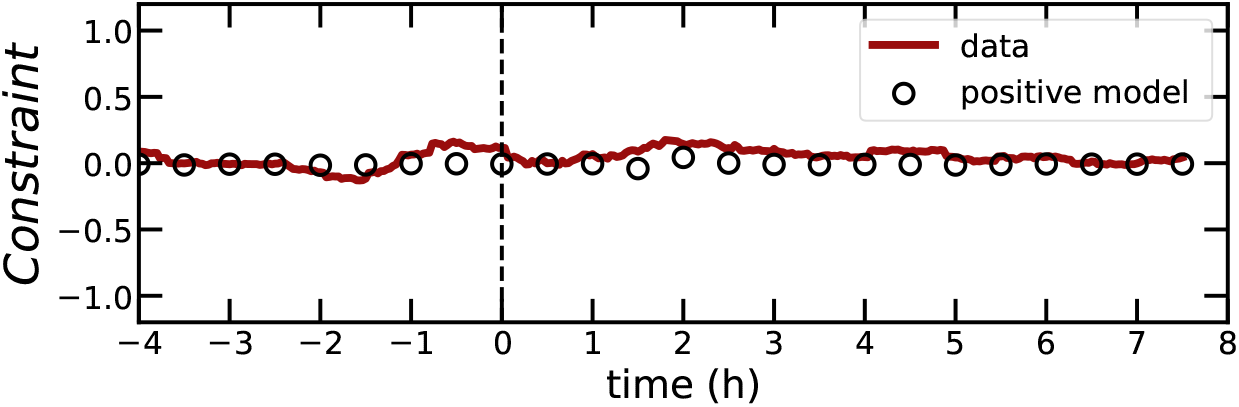
Both data and model fulfill the expected constraint *(e^G^*^(1^*^−λ^*^)^*/*(1 + *ζ*)〉 = 1, where *G* = log(*V_f_ /V*_0_), *λ* is the slope of the size-growth plot and *ζ* is the slope of the adder plot. This constraint holds theoretically beyond steady state, and the data confirm this prediction. The constraint can be derived from the definition *ζ* = *d*Δ*/dV*_0_ noting that, if *q* = log(*V*_0_), by chain rule *d*Δ*/dq* = (*d*Δ*/dV*_0_)(*dV*_0_*/dq*) = *ζe^q^*, and equally, since Δ = *V_f_ − V*_0_, *d*Δ*/dq* = *e^q^ e^G^*(1 + *λ*). The verification of this constraint provides a useful consistency check for our analysis.

**FIG. S6.**
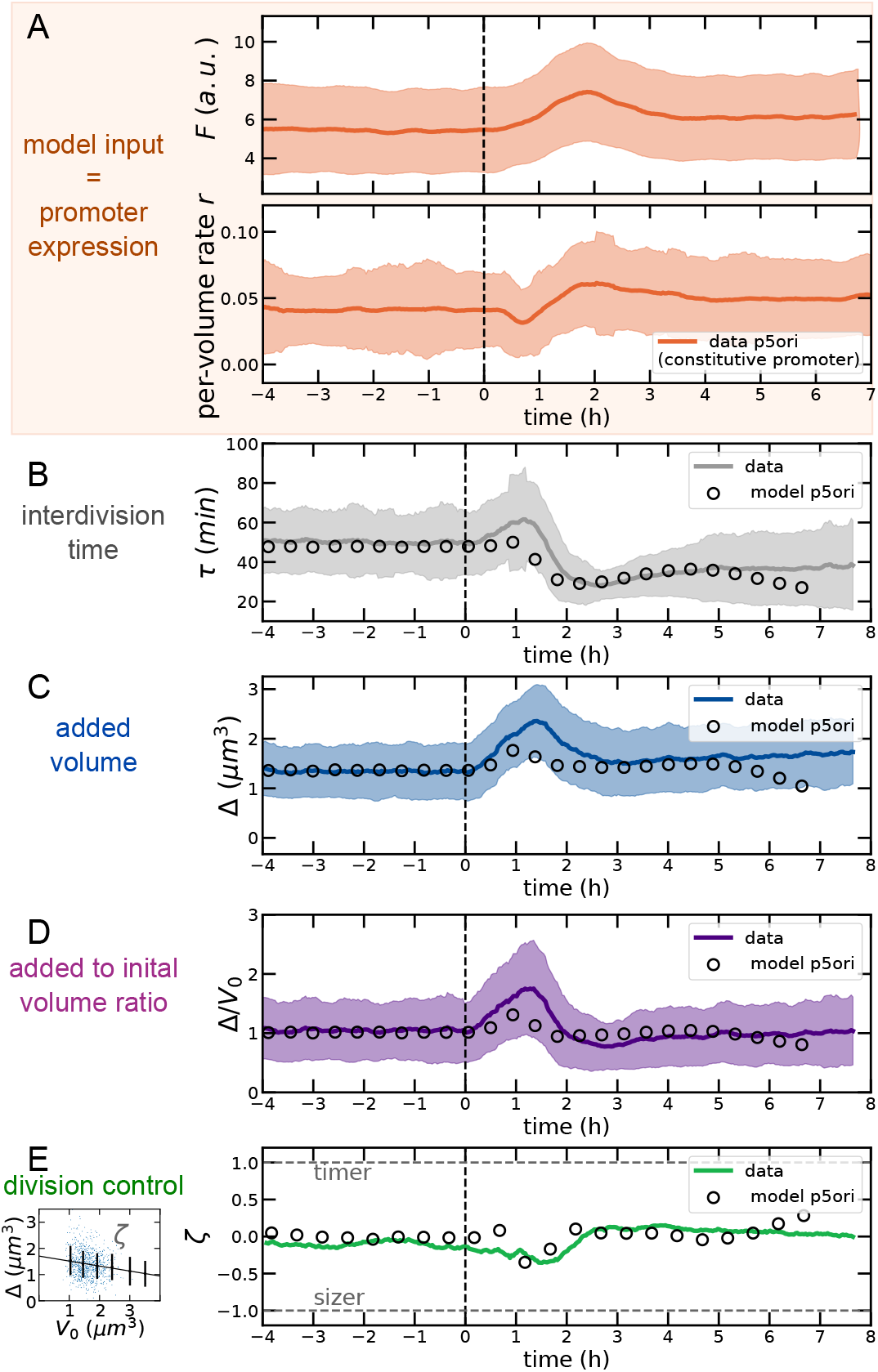
Results of the divisor protein model using as input the measured volume-specific production rate *r* from the *P5 constitutive promoter* inserted close to the *replication origin*. All panels as in Figure 4 of the main text.

**FIG. S7.**
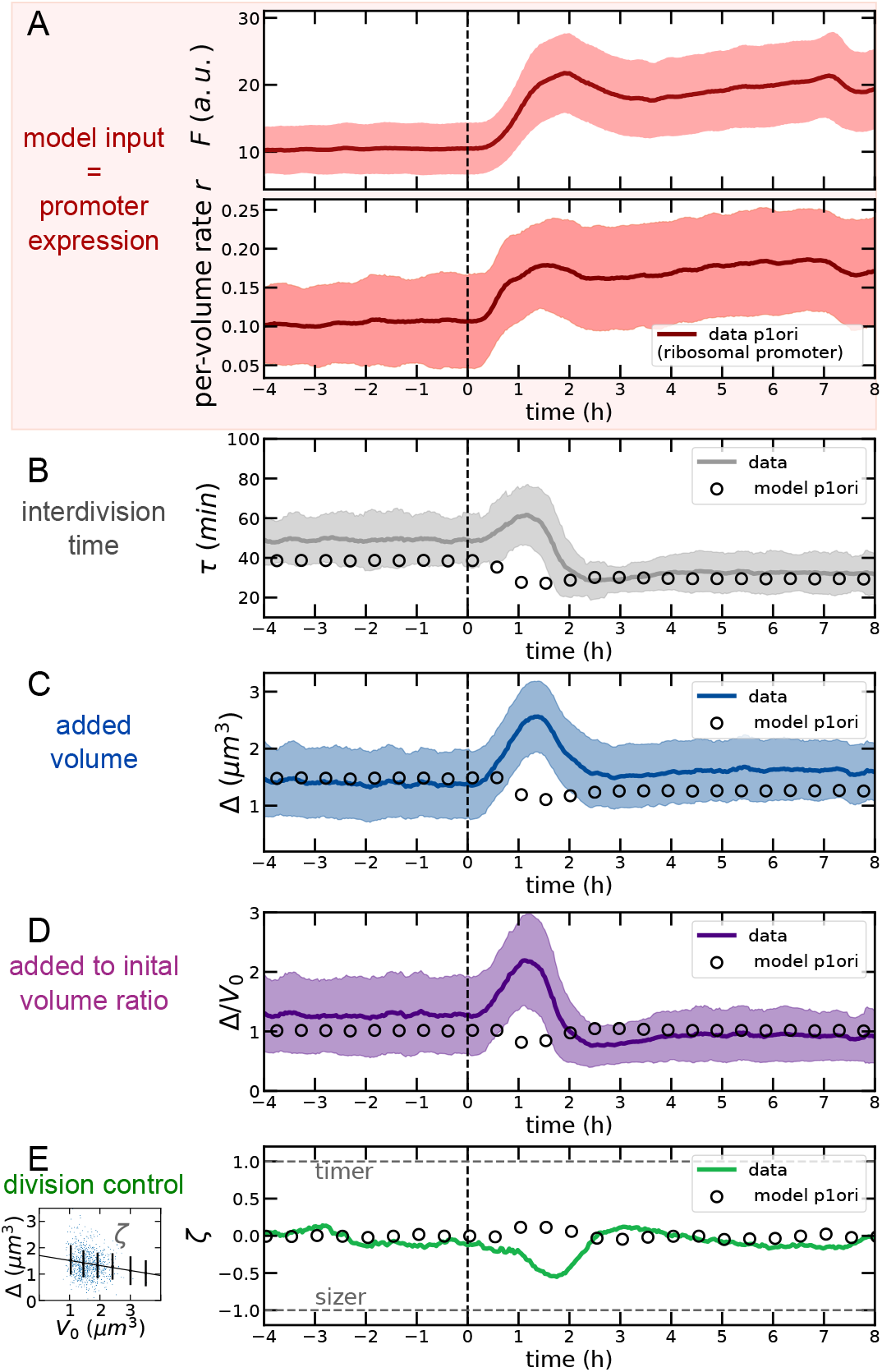
Results of the divisor protein model using as input the measured volume-specific production rate *r* from the *P1 ribosomal promoter* inserted close to the *replication origin*. All panels as in Figure 4 of the main text.

**FIG. S8.**
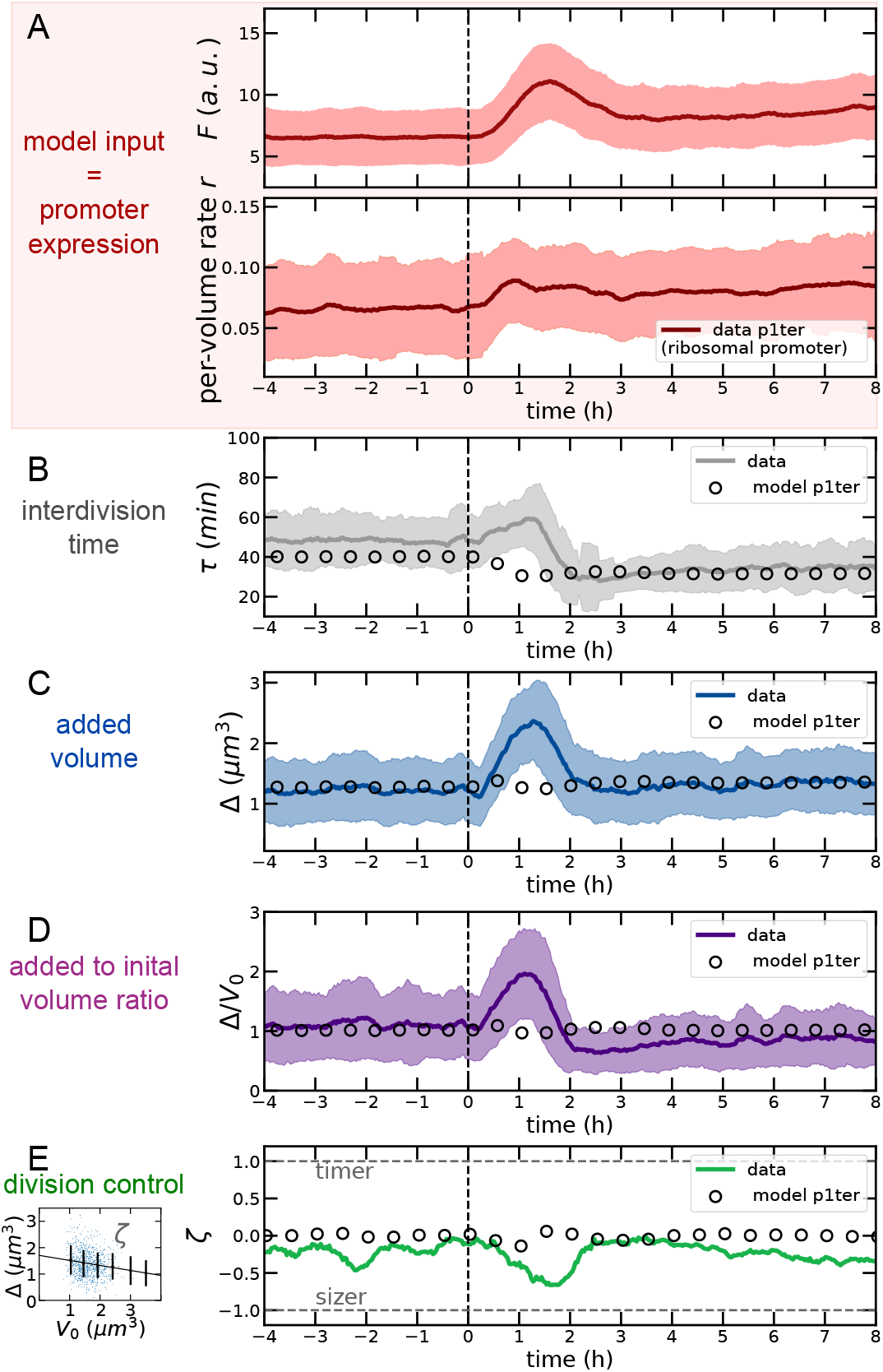
Results of the divisor protein model using as input the measured volume-specific production rate *r* from the *P1 ribosomal promoter* inserted close to the *replication terminus*. All panels as in Figure 4 of the main text.

**FIG. S9.**
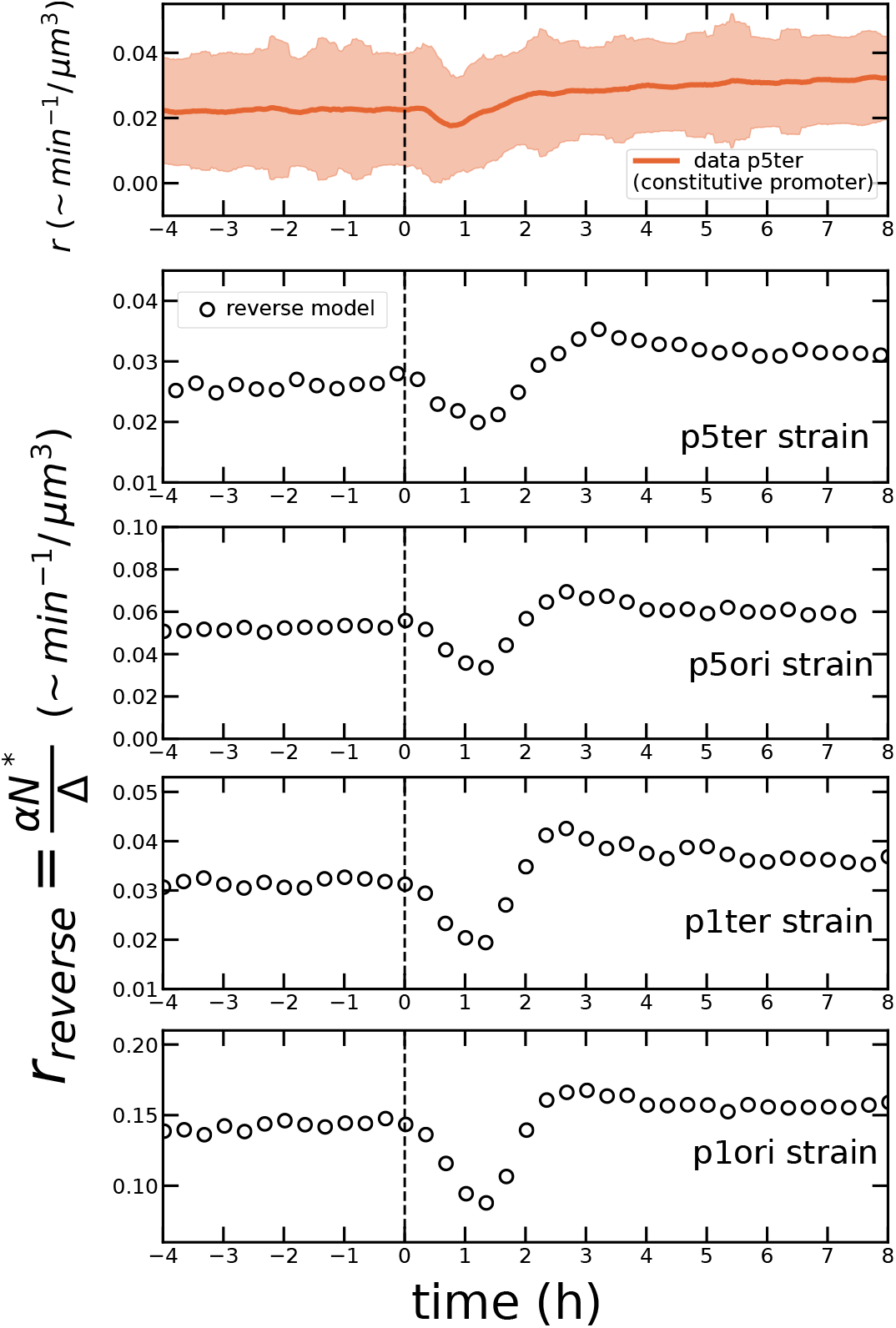
The theoretical volume-specific production rate of the adder molecule derived from data on growth rate and added size transiently decreases after the shift, before reaching a higher plateau. The top panel is the measured volume-specific rate of GFP from the p5ter promoter shown in Fig. 4 of the main text. The other plots are binned averages of 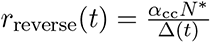, shown in the different panels to be robust across shift experiments with strains carrying different promoters. The reverse argument assumes that *N^∗^* is constant after the shift. In these plots, *N^∗^* is set to the value of 2 (in arbitrary units, taken from Supplementary Figure S4) before and after the shift, in order to make the model match the data also quantitatively. The average uses experimental values of Δ = *V_f_ − V*_0_ and of the growth rate *α*_cc_ comes from exponential fits of the volume *vs* time data of each cell cycle, and the variables are associated to the division time of a cell.

**FIG. S10.**
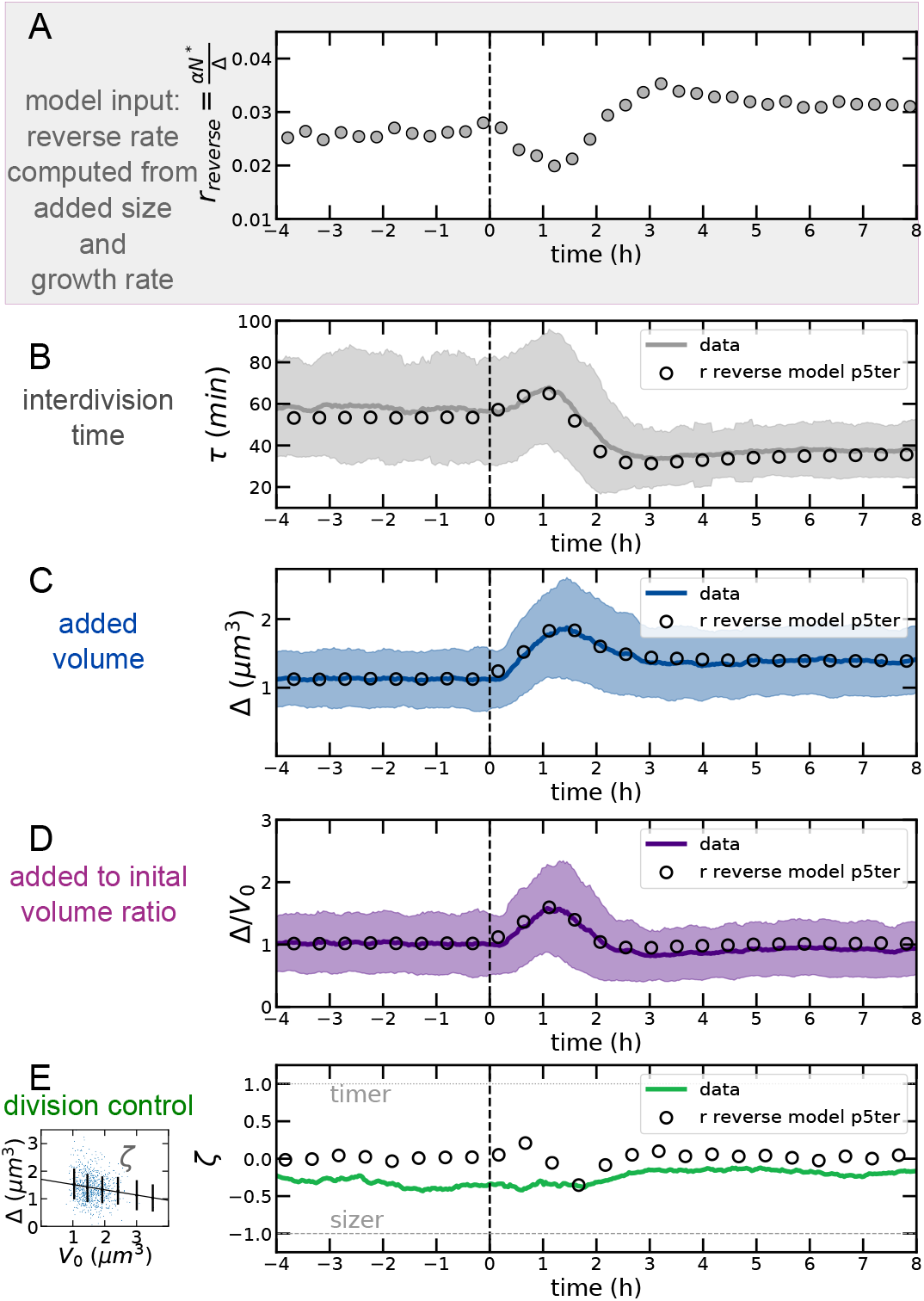
Results of the divisor protein model using the reverse production rate *r*_reverse_(*t*) inferred as described in Fig. S9 using the strain carrying the P5 constitutive promoter close to the replication terminus. All panels as in Figure 4 of the main text.

**FIG. S11.**
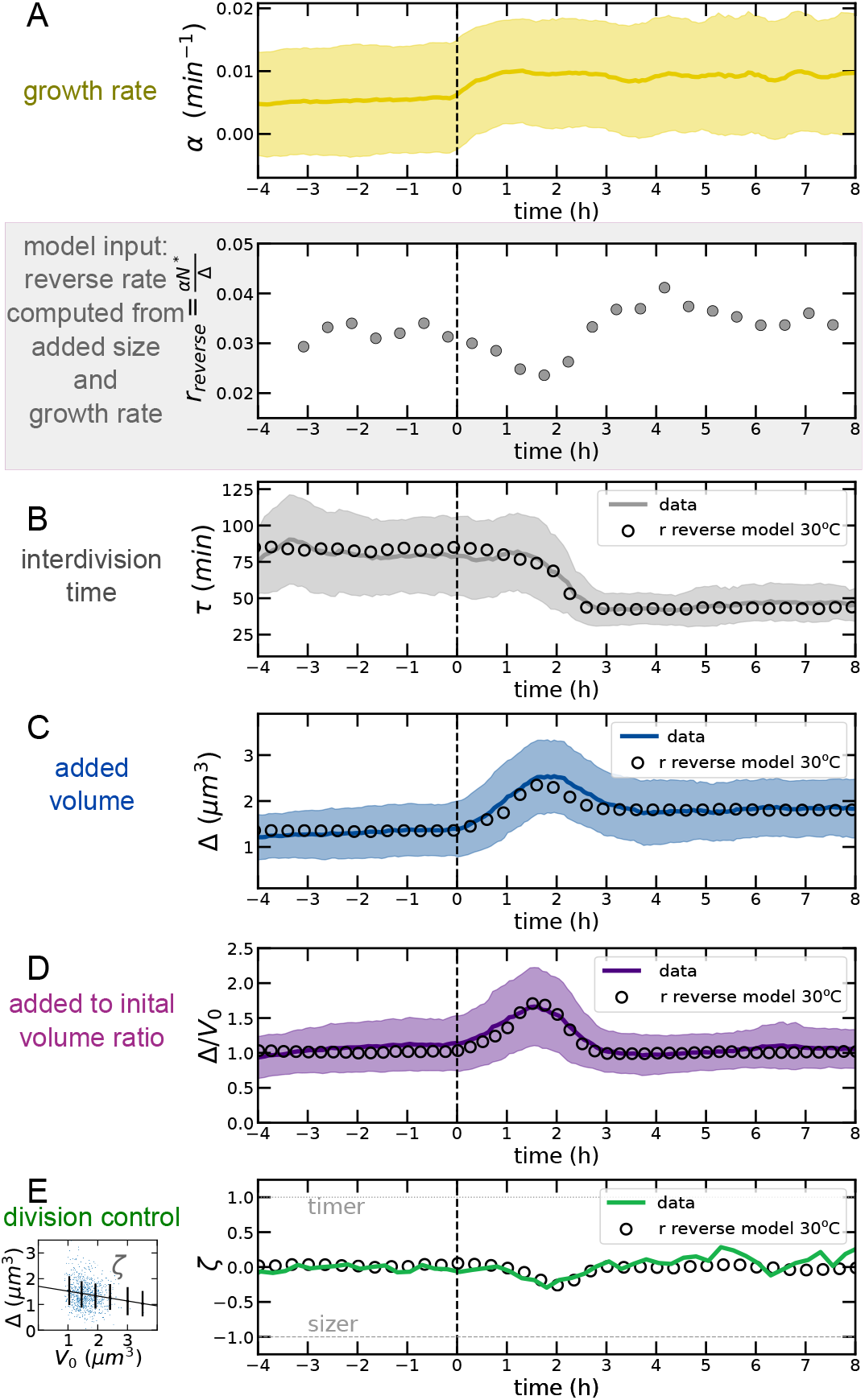
Results of the divisor protein model using the reverse production rate *r*_reverse_(*t*) for an experiment at 30*°*C using the strain carrying the P1 constitutive promoter close to the replication origin. All panels as in Figure 4 of the main text, except for panel A, which also includes the growth rate for this different experiment.

**FIG. S12.**
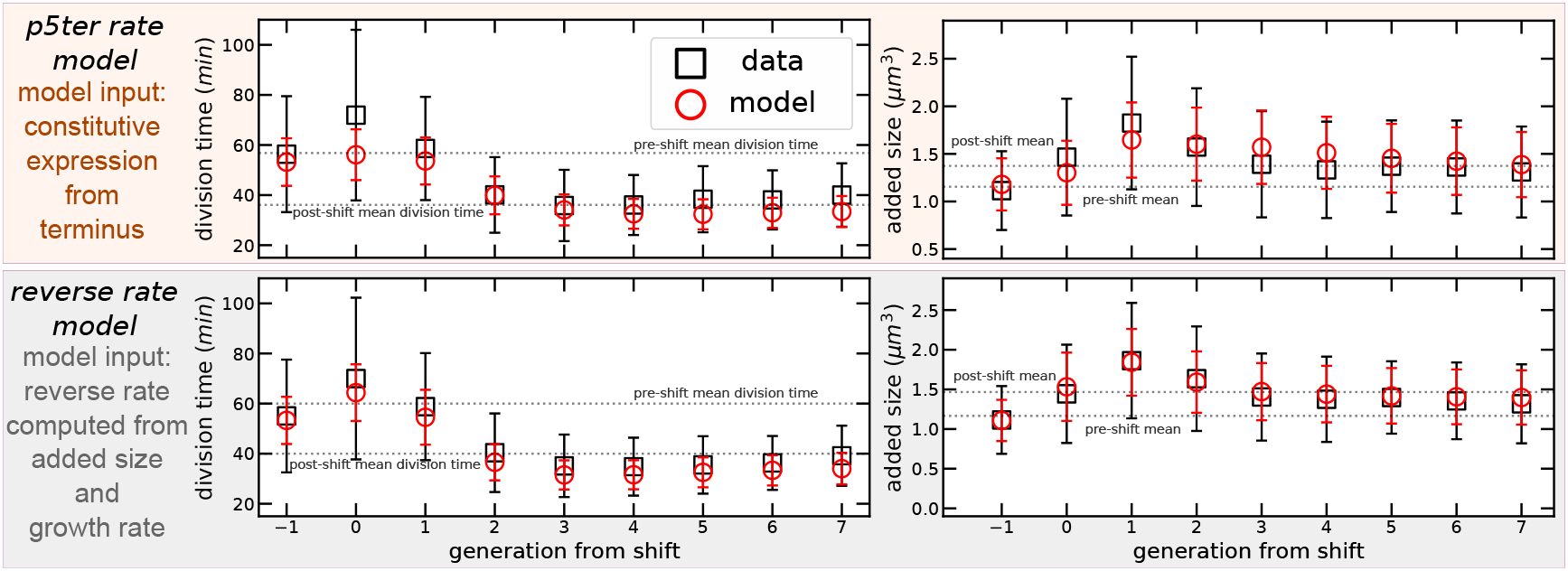
Cell-division behavior by generation from/to the nutrient shift. The generation index is set to 0 for the cells that see the shift during their cell cycle. The plots compare data (circles) with the forward model (top) using P5ter promoter production (see Fig. 4 in the main text) and with the reverse model (bottom, see Fig. S9 and S10). Both interdivision time (left panels) and added size (right panels) show overshoots (equivalent to the ones seen in Fig. 2 of the main text) that are reproduced by the model. Error bars are standard deviations

**FIG. S13.**
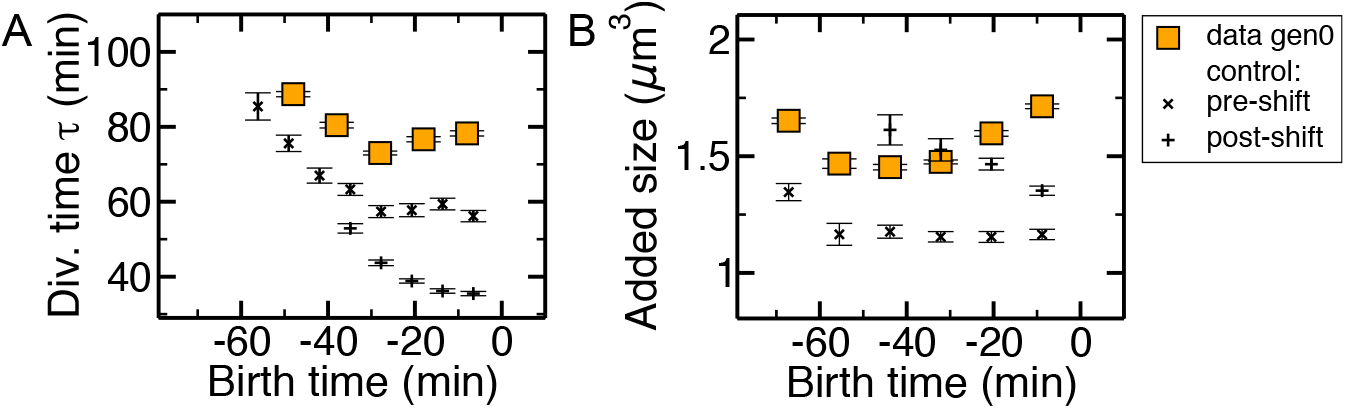
The behavior of the generation seeing the shifts differs from the pre- and post-shift background in a cell-cycle dependent way. The plots report mean interdivision time (A) and added size (B) as a function of birth time (time 0 is the shift) for “generation 0” cells, which see the nutrient shift during their cell cycle. Control data report the same quantity for cells in the steady pre-shift (crosses) and post-shift condition, wuth respect to an arbitrary reference time. Error bars are standard errors.

**FIG. S14.**
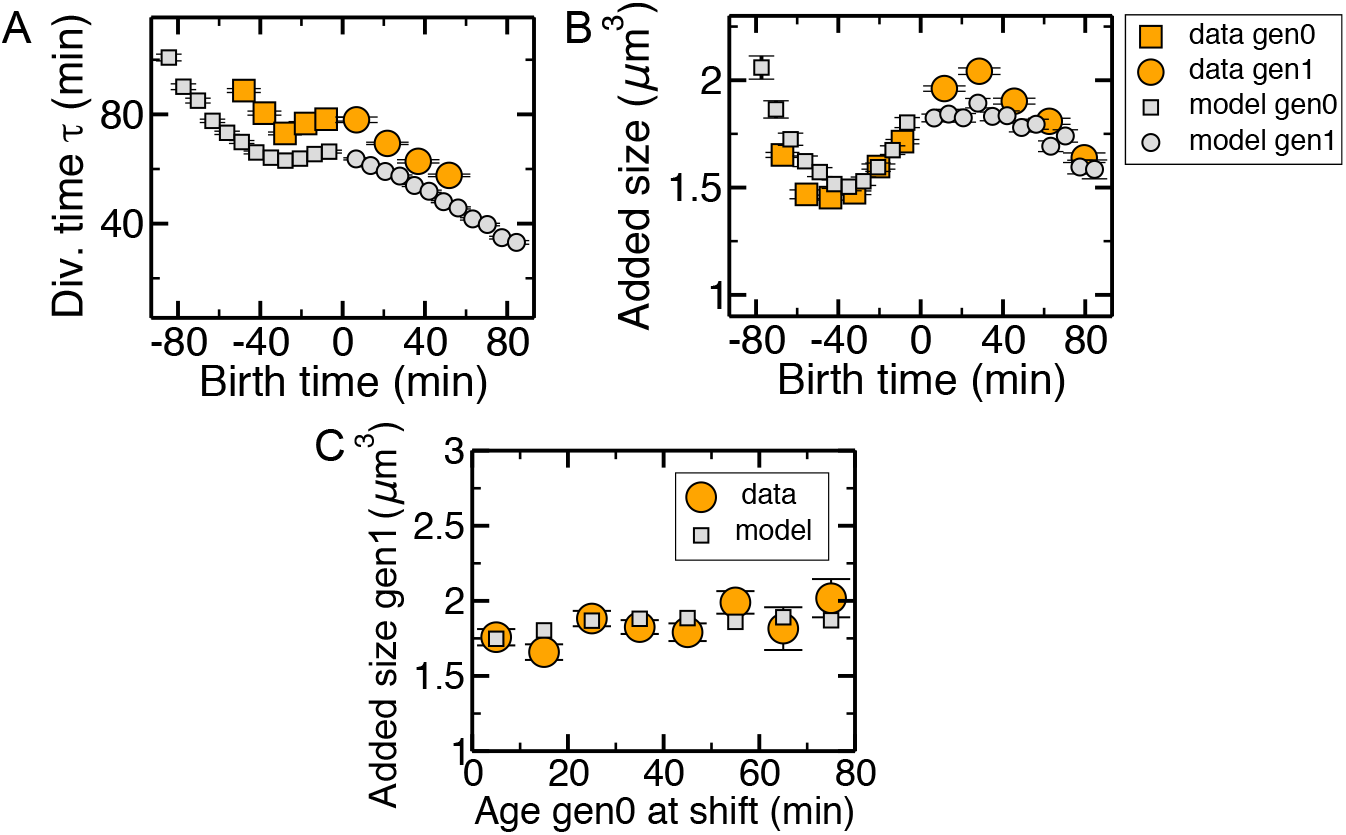
The behavior of the generation seeing the shift and the one after is reproduced qualitatively by the accumulator model. The top panels report interdivision time (A) and added size (B) for for “generation 0” (orange squares) cells, which see the nutrient shift during their cell cycle and for their daughters, “generation 1” cells (orange circles). Grey squares and circels show the equivalent results for the reverse model. C) The added size of generation 1 cells does not depend on the cell-cycle time at which their mothers saw the shift, in both model and data. Error bars are standard errors.

**FIG. S15.**
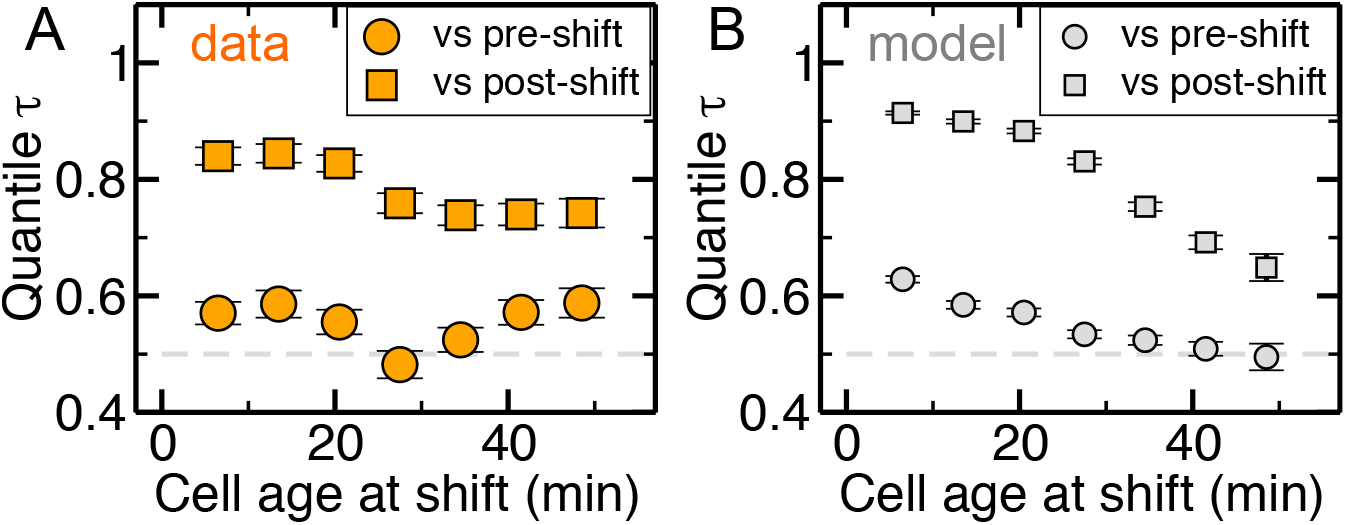
The cell-cycle pattern in division delays differs between model and data, suggesting that the checkpoint is also related to additional processes not described by the model, such as chromosome replication and segregation. The plots report the mean quantile of the interdivision time for “generation 0” cells, which see the nutrient shift during their cell cycle, as a function of the time from firth when they see the shift. The *y* axis quantifies the position of the cell cycle time of a cell compared to the cells of its same age that do not see the shift. A quantile of 0.5 means that the cell behaves as pre- or post-shift cells. In the model, cells that see the nutrient shift late in their cell cycle do not modify their cycle duration compared to the pre-shift conditions, while in the data these cells delay their division. Error bars are standard errors.

**FIG. S16.**
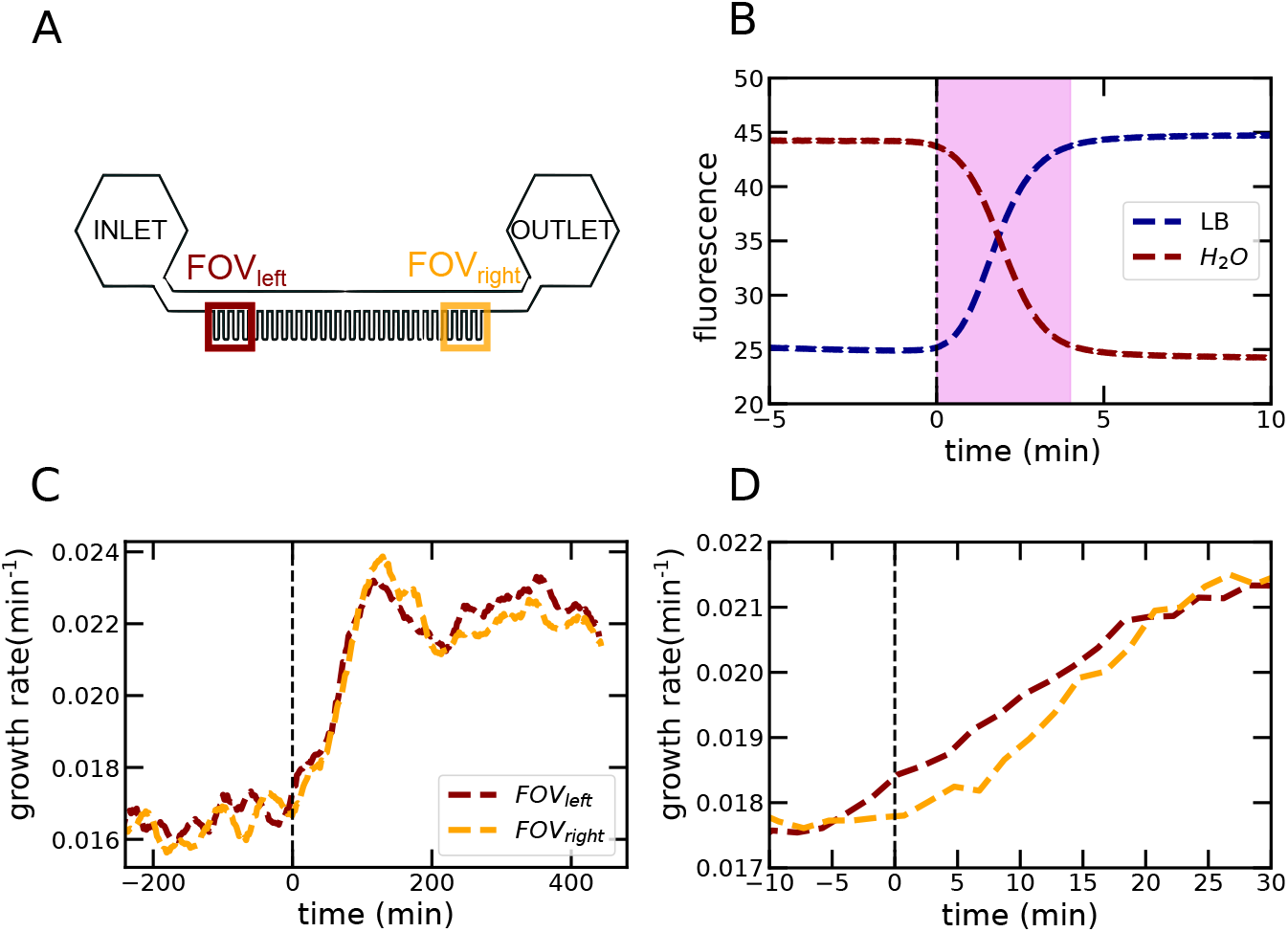
Test of filling time and delays in nutrient availability within the microfluidic device. A) Scheme of the device with the outermost fields of view labelled in dark red and orange. B) Experiment loading the device with fluorescent LB medium and monitoring mean fluorescence within the channels. Side channels start seeing the new medum immediately, and completely fill in about 4 minutes (purple shaded area) once the medium enters the main channel (*t* = 0). C) The delay between growth rate changes in the experiment averaged in the outermost fields of view in the device is small. D) Zoom of panel C showing that the delay is about 5 minutes.

## SUPPLEMENTARY INFORMATION ON THE FRAMEWORK FROM CADART *ET AL.* 2018

This appendix describes the framework from refs. [15, 17] used in Supplementary Fig. S2. This analysis assumes a random exponential growth rate that may be dependent on a cell’s initial size. The quantity *G* = log(*V_f_ /V*_0_) is the overall multiplicative growth. Size control here is quantified as the linear correction to deviations from the mean logarithmic size (i.e., smaller-born cells grow more than larger-born ones). Three key parameters indicating the degree of size control fall from the procedure [15, 17]. The overall size control is 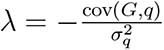, where *q* = log *V*_0_. The value of *λ* signals the size control strategy in use. Specifically, *λ* = 1 corresponds to a sizer (constant target size); *λ* = 0 represents no control; and *λ* = 0.5 denotes an adder. The growth and timing components of size control are, respectively, 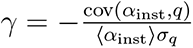 and 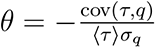. A positive value for either *γ* or *θ* indicates a homeostatic effect while negative values mark a negative contribution to size control. for example, for positive *θ*, smaller-born cells grow for longer periods of time, and for positive *γ* they grow at a faster rate. Viceversa, for negative *γ*, growth rate contributes to variability in the initial size. The three parameters yield a quantitative method to determine the contribution of timing and growth-rate corrections to size homeostasis. They are mathematically related through the expression

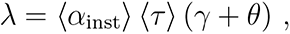

which we find verified in the data (Fig. S2C). At steady state, this analysis confirms that *E. coli* division control is adder-like and that this strategy is realized primarily through timing control, while fluctuations in growth rate have little influence (Fig. S2A). This is consistent with previous analyses [15, 17].

Departure from the steady state occurs 20 min after the shift, when the average added volume exceeds the average birth volume. This ratio is maximized at 1.1 h after the upshift at 1.64 : 1, equilibrating back to 1 : 1 after about 4 h (Fig. S2C). The shape of this response is largely expected because we have an approximately closed system in the sense that nascent cell lineages are not added to the device after the initial incubation and trapping, hence individual birth volumes cannot increase without an increase of added volume in the previous cell cycle. This result is not trivial though: it is conceivable that a perturbation could briefly arrest cells that were smaller than average at birth, prompting larger cells to overtake the dividing population and thereby increasing birth volume without a preceding increase in added volume. Despite the necessary break from the steady state (Δ*) / (V*_0_〉 = 1, the overall strength of size control is nearly constant during the upshift as evidenced by *λ*, which exhibits only a moderate monotonic decrease from 0.638 *±* 0.046 in slow growth media to 0.516 *±* 0.033 in fast media (Supplementary Fig. S2CD). Thus, although there is a change in the overall added volume, cells demonstrate near-adder behaviour that is uninterrupted by the environmental shift.

This persistence is effected by complementary dynamics between the timing- and growth-related components of size control. The nutritional upshift induces a brief but dramatic decrease in the normalized growth-related contribution 〈*α*_inst_*〉 (τ) γ*, which pulses downwards from a negligible *−*0.019 *±* 0.101 in the first steady state to a minimum of *−*0.466 *±* 0.298 after 1.6 h in the second media (Supplementary Fig. S2CD). Growth-related control then returns to a comparatively small equilibrium value of *−*0.176 *±* 0.090. Control from timing, 〈*α*_inst_*〉 (τ) θ*, responds in an opposing manner, increasing considerably from 0.551 *±* 0.335 preshift to a 1.107 *±* 0.679 maximum 1.5 h into the upshift (Supplementary Fig. S2CD). It too equilibrates after about 3 h, settling to 0.665 *±* 0.310. Therefore, although growth and timing processes are thrust out of their respective equilibria by the media shift, the balanced interplay of their contributions to cell size ultimately conserves near-adder behaviour in this dynamic environment (Supplementary Fig. S2D). It should be noted that *λ* = 0.5 strictly coincides with a perfect adder provided it is at steady state (Δ*) / (V*_0_〉 = 1. Generally, the birth size-independent addition of an arbitrary volume during the cell cycle that defines an adder can present a range of *λ* values: e.g. if the added volume is between 0.5 and 2 times the average birth volume, *λ* would be in the range [0.33,0.67]. Nevertheless, *E. coli* is well within the adder regime at all time points.

## SUPPLEMENTARY INFORMATION ON THE *BUGPIPE* SEGMENTATION/-TRACKING ALGORITHM

Bugpipe is the custom-coded MATLAB-based package that was used to segment, track, and analyse the resultant cell data presented in this work. The following contains a summary of the algorithms developed. The package and its full documentation is available at https://github.com/panlilio/bugpipe.

### RAW IMAGES USED

Raw data consisted of 16-bit images at 512 x 512 pixel resolution. Three images were taken per field of view, per time point: one fluorescence; one brightfield; and one dark frame for background subtraction, i.e. an image acquired under the same exposure time but without illumination (Fig. S17). Images were obtained every 5-6 min for each field of view, with the number of fields chosen for continuous acquisition. This translated to a total of 35-40 fields of view, each spanning up to 8 microchannels. The chip was aligned such that the microchannels run approximately parallel to the image *x*-axis.

Fluorescence images were used for segmentation here because of the associated high signal-to-noise ratio. In principle, high quality phase contrast images can be successfully substituted with minor changes to the segmentation procedure. Brightfield images were used largely for manual inspection when required.

**FIG. S17.**
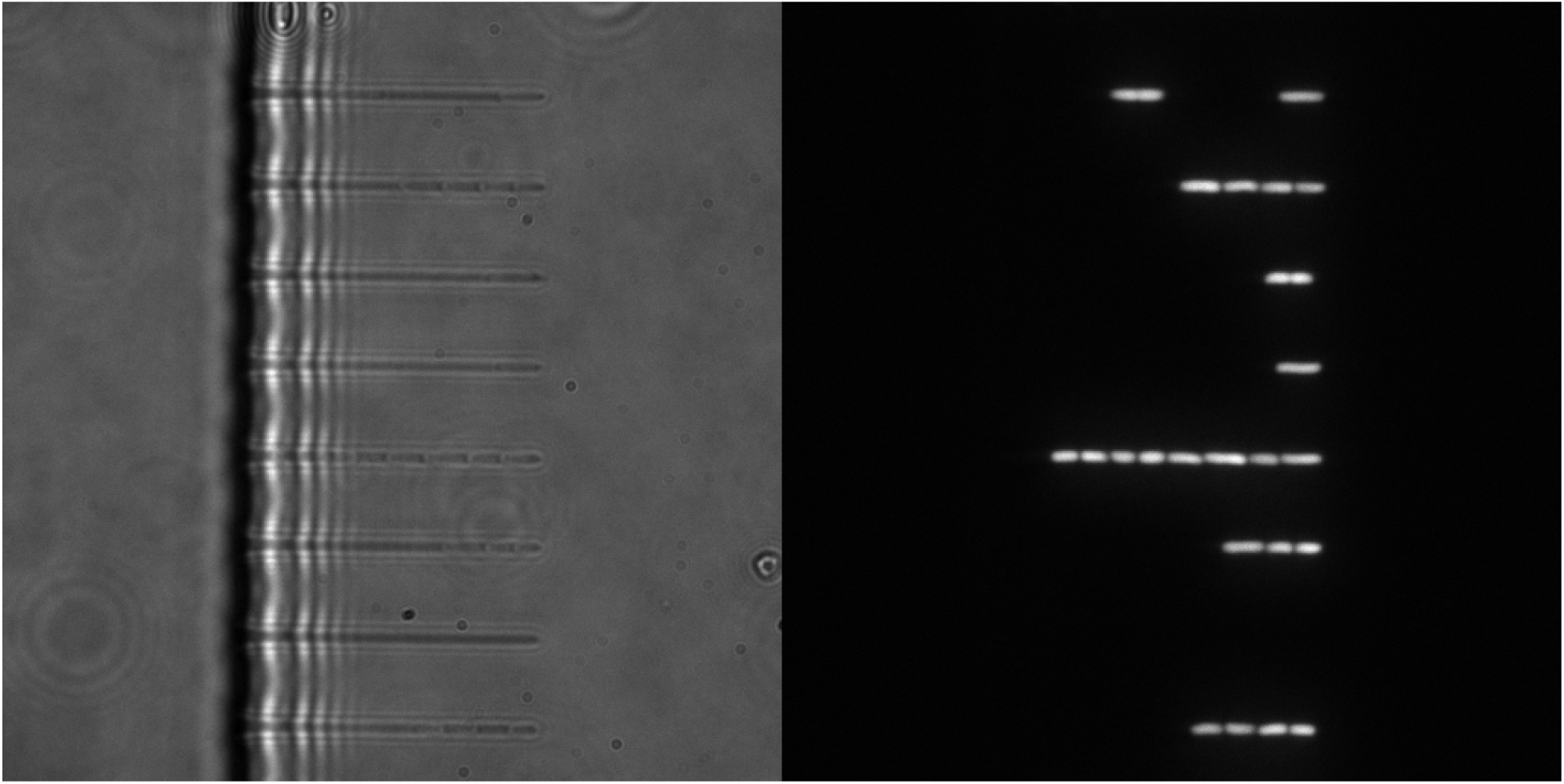
Sample brightfield (left) and fluorescence images (right).

### BUGPIPE: IMAGE SEGMENTATION

Initially, all images are segmented independent of the order in which they were acquired by the function `segmentationMP`. For each background-subtracted fluorescence image the algorithm proceeds in the following steps: 1) find regions of interest, i.e. the micropistons trapping each cell line; 2) within each region of interest, threshold and perform morphological operations to isolate cells in the background; 3) find cell boundaries and separate any artifically fused cells; 4) remove artefacts based on size.

The order-independent first pass at segmentation enables parallel processing of all frames and thereby imparts a drastic decrease in the overall characterization time. Corrections to the initial segmentation are performed under the tracking algorithm. All programming was done in MATLAB with built-in functions indicated by (*^†^*).

#### Background subtraction

Simple background subtraction is performed by subtracting the respective dark frame from each fluorescent image. Any resulting negative intensity values are set to 0.

#### Microchannel detection

The regions of interest (ROI) in each image are defined by the microchannels trapping each line of cells. The function `getChannelsMP` uses *x*- and *y*-cross sections of pixel intensity to detect these regions by thresholding: namely, the Otsu threshold is used along *x* (parallel to direction of growth, Fig. S18A, green) and a sliding average background subtraction is implemented along *y* (perpendicular to growth, Fig. S18A, blue). The final regions of interest are shown in Fig. S18B. The typical microchannel region delineated by this procedure across all fields of view and replicates was around 35 x 200 pix.

#### Cell segmentation

Each region of interest is processed independently. The following procedure is implemented by `segmentationMP` as outlined in Fig. S19A. First, the cropped image of the *k*-th microchannel *I^k^* is passed through a 2D median filter to remove hot pixels (`medfilt2`*^†^*) and then resized using bilinear interpolation (`imresize`*^†^*, 3x magnification) to further smooth intensity values before thresholding.

**FIG. S18.**
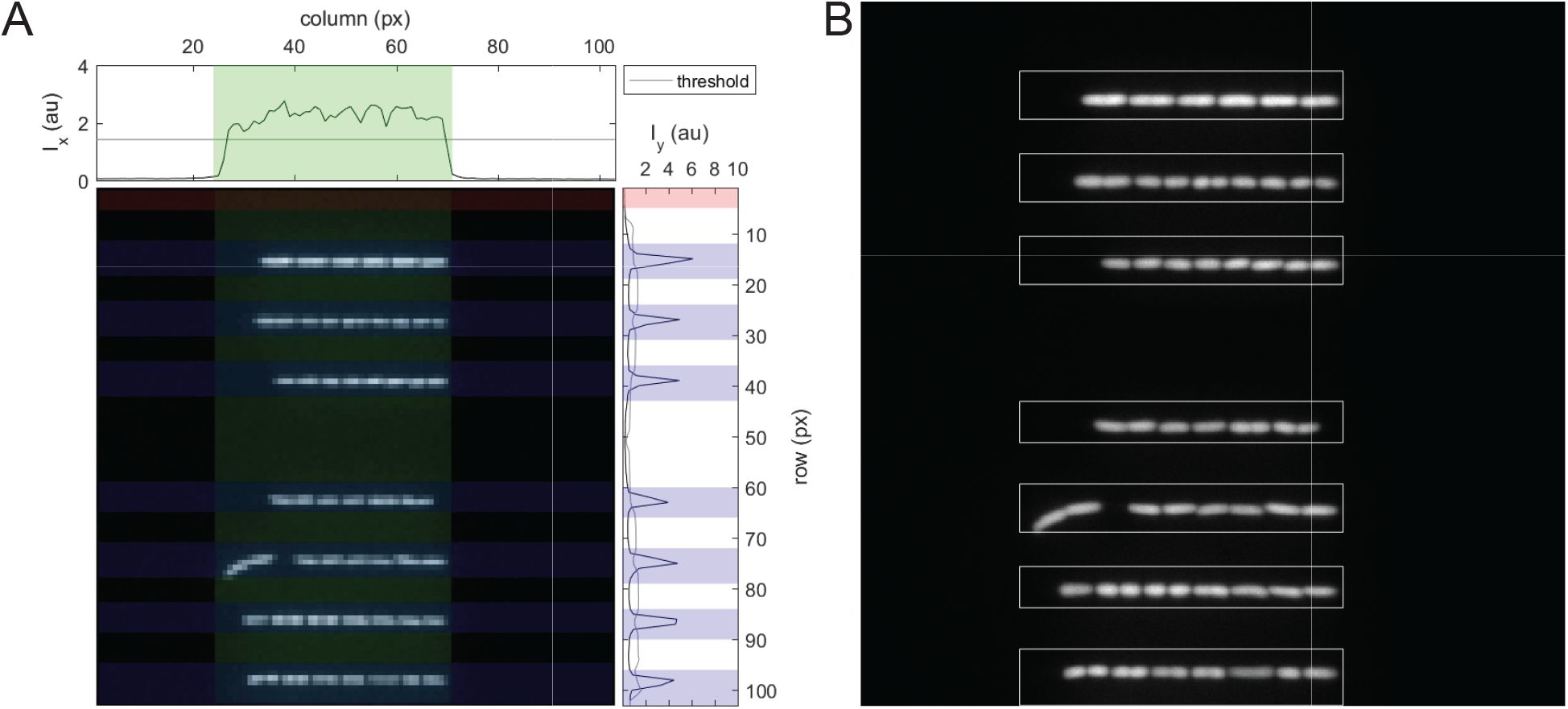
Microchannel identification by thresholding image cross sections of intensity **A** Subsampled image where the *x*-profile *ι_x_* defines the height of all channels in a given field of view (green) and *ι_y_* determines the width of each microchannel (blue). Regions that pass the filter but do not contain any high intensity objects are omitted from further analysis (red). **B** Found microchannel bounding boxes overlayed on image at its original resolution.

#### Modified Otsu threshold

The cropped greyscale image *I^k^* is thresholded using Otsu’s method with one modification (Fig. S19B). The standard Otsu threshold (as calculated by `graythresh`*^†^*) tended to drastically undersegment the fluorescence images here: i.e. several cells or sometimes entire microchannels were grouped into a single object, often lacking a sufficiently detailed perimeter from which cells might be distinguished. To overcome this, a correction term was added equal to the standard deviation of the Otsu-determined lower intensity population. That is, if *T*_Otsu_ is the Otsu threshold and the set of low intensity values is 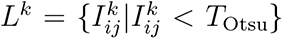, the applied threshold is

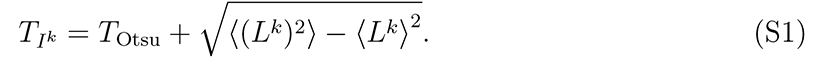

All 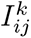 less than *T_I_k* are set to 0 and all pixels greater or equal to the threshold are set to 1. The result is a binarized image of black 0’s and white 1’s (Fig. S19A, 3rd panel down).

The thresholded image is then morphologically dilated using a 3 pixel radius disc (`imdilate`*^†^*) to compensate for the slight oversegmentation during thresholding and further smooth object edges. The boundaries of the resulting objects are then found with `bwboundaries`*^†^*.

#### Curvature assessment

Each object was then assessed by the function `getCellsMP` according to its ordered (*x, y*) boundary points, with each coordinate pair representing an edge pixel. For a single bacterium this set typically consisted of about 120 points on the magnified image. Although a large fraction of detected objects were indeed individual bacterium, the above procedures did tend to artificially join multiple cells by undersegmentation. In this case, the boundary of the joint object consistently had at least two regions of high magniture, negative curvature near the point of contact between adjacent cells (Fig. S19C).

For a single rod-shaped cell, curvature is theoretically non-negative for the entire boundary. Candidate objects for further segmentation are therefore identified as those possessing two or more regions in which the above smoothed curvature is less than −0.03 pix*^−^*^1^. This gives an allowance for some noise in the image and in practice serves as an empirical threshold for separating emerging daughter cells during cell division.

Objects without multiple negative curvature regions were rescaled to match the original 512 x 512 pix image and assessed for area using `regionprops`*^†^*: to remove the rare imaging artifact, only those objects consisting of more than 30 pix (*∼* 0.034 *µ*m^2^) were accepted as individual bacterium. Note that a typical bacterium has a corresponding projected area of roughly 156 pix (*∼* 1.79 *µ*m^2^). The aforementioned candidates for further segmentation were passed to the next processing step.

#### Boundary processing

The local minima of negative curvature regions are each designated as a “pinch point” and, owing to the configuration of the cells, generally one pair of pinch points can be connected to separate two adjacent cells. That is, because cells tend to stack along their major axes, two pinch points appeared at the point of contact between two cells: one above and one below the axes (Fig. S19C, second and third panels down). In the case of more than two pinch points, it was neccesary to calculate each point’s nearest neighbour for correct pairing. In many cases, only two bacteria are enclosed in the object and they are easily separated using a pair of pinch points.

For example, consider the case of two merged bacteria. Line segments were demarcated by pinch points. Each segment was then closed by connecting its first and last points. This connecting line was interpolated at *n_c_* points using the first and last *n_c_* points of each segment, assuming a third degree polynomial, where *n_c_* was the approximate number of pixels along the straight line between the endpoints. The closed segments were then rescaled to match the original 512 x 512 pix image and passed by the same size filter to ensure that the area enclosed was greater than 30 pix. The final cells segmented for the sample channel are shown in the bottom panel of Fig. S19A.

**FIG. S19.**
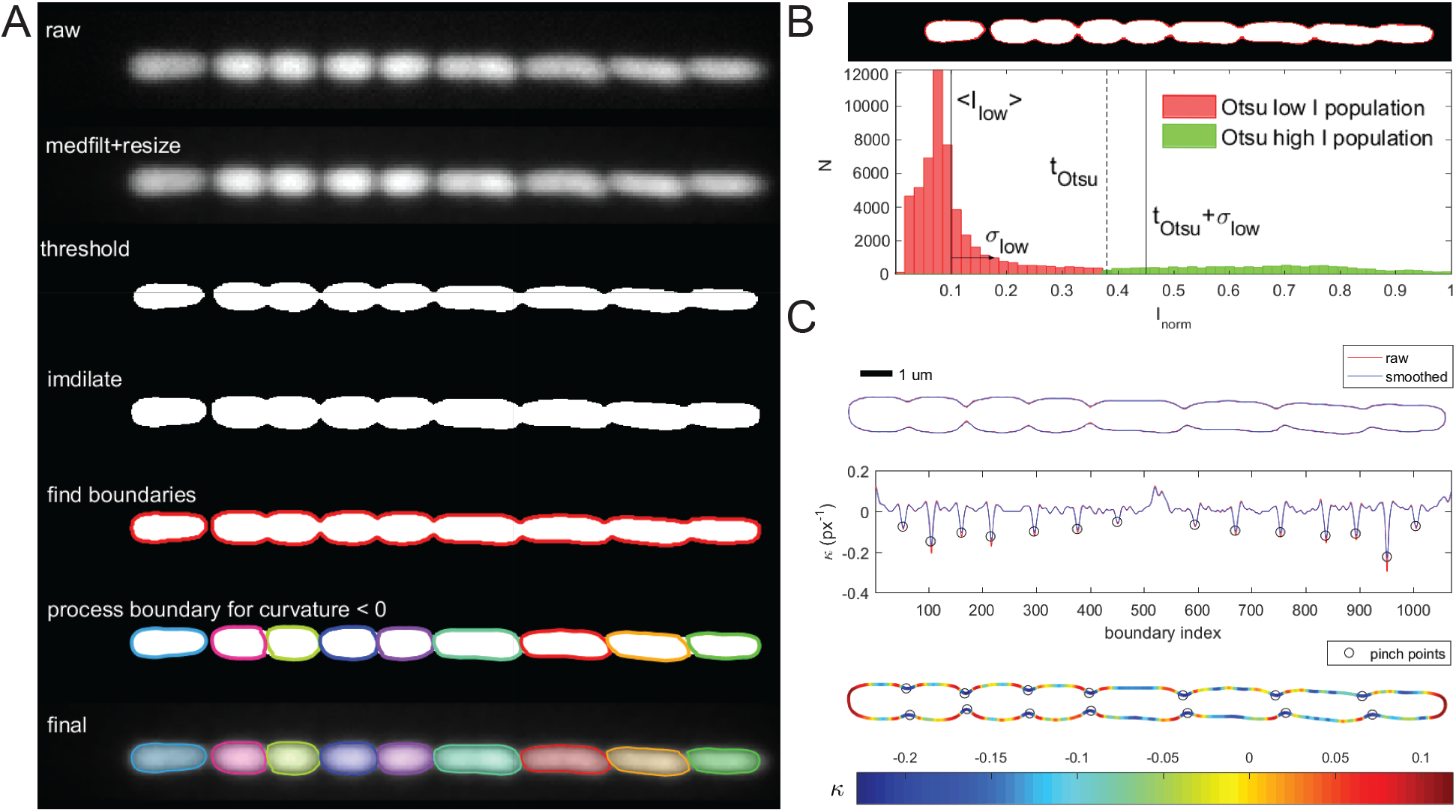
Morphological operations and object boundary processing for cell segmentation within a microchannel. **A** Top to bottom: overview of the full segmentation process on a single microchannel. **B** Demonstration of the modified Otsu threshold: histogram of mean-normalized pixel intensities for the channel of interest, with the red population indicating the background (i.e. low intensity) pixels and green indicating object (high intensity) pixels according to the standard Otsu threshold (dotted vertical line). Difference between the Otsu threshold and modified threshold is shown in upper panel (red). **C** Segmentation of the large, merged object in A using boundary curvature (second panel). Boundary coloured according to local curvature (third panel) with regions of strongly negative curvature (dark blue) indicating pinch points at which to further segment cells (black circles).

This approach based on boundary curvature turns the problem of segmentation from 2D to 1D and minimizes the use of the repeated morphological operations often required for e.g. watershed-based separation. Rather than smoothing cell edges by morphological opening and/or closing, images are instead slightly overthresholded then dilated in order to retain the regions of sharp negative curvature anticipated at the fusion points—real, during the process of cell fission, or artifactual—between two cells. The resulting segmentation yields results comparable to that produced by edge detection filters but avoids the more time-consuming morphological operations.

### BUGPIPE: CELL TRACKING

An order-dependent approach is blatantly necessary for frame-to-frame cell tracking. The time series for separate fields of view are independent however and subsequently they are processed by separate MATLAB workers (`spmd`*^†^* in the Parallel Computing Toolbox). The tracking procedure takes advantage of the geometric constraints imposed by the mother machine’s format: in particular, that in the absence of cell divisions a cell’s rank within a channel is conserved between frames. That is, suppose one channel has five cells in one frame, five in the next, and there was no evidence of cell division. The first cell away from the dead-end of the channel in one frame would be the first cell away from the dead-end in the next frame, the second cell in one frame would be the second cell in the next frame, and so forth. The task was thus reduced to: 1) microchannel tracking between frames, 2) sorting cells in-channel based on distance from the dead-end, and 3) determining markers for cell division to adjust the rank-based pairing as necessary. These operations are all contained in the function `cellTrackerMP`, which takes consecutive images and the results of their respective segmentations as input.

#### Microchannel matching

The location of each microchannel’s bounding box is passed through the segmentation variables associated with each frame. Pairwise-distance calculations are perfomed for channel tracking because of the small channel numbers: there are always fewer than 9 microchannels under the reported magnification and camera conditions. This approach assumes sufficiently low optical drift, specifically that any relative movement between consecutive frames is less than the half-distance between microchannels: a criterion that was easily met by the previously described microscopy system.

Example matched channels are shown in Fig. S20.

**FIG. S20.**
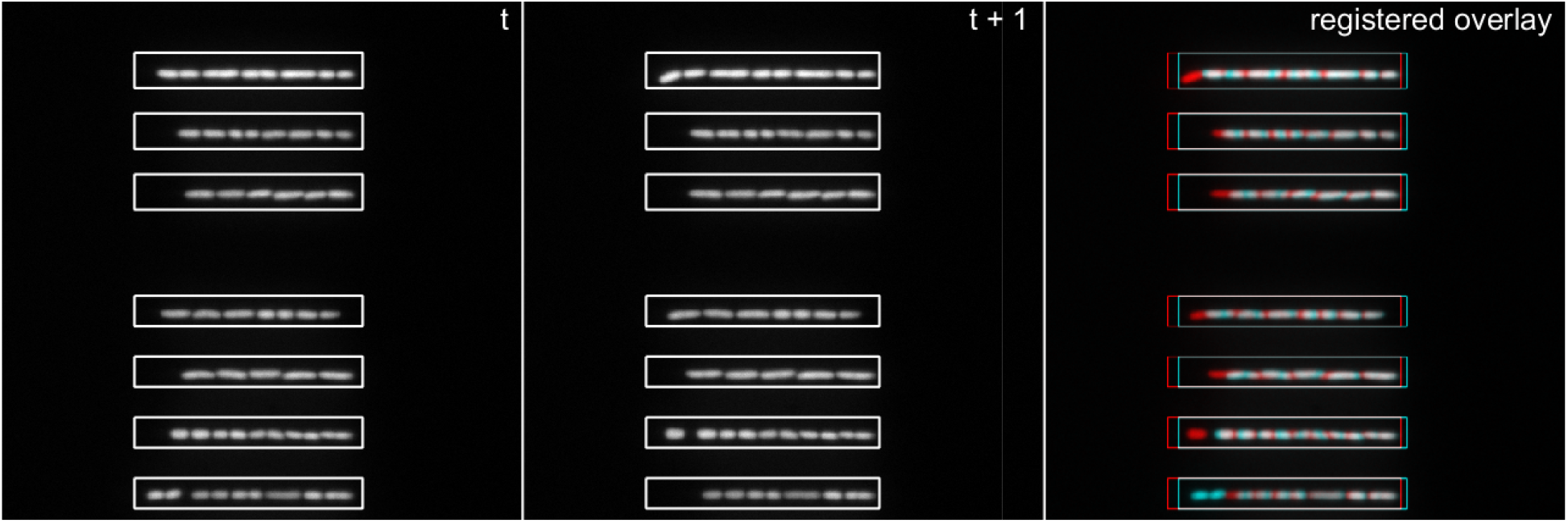
Microchannel detection and pairing between frames. Bounding boxes identified for the microchannels in two consecutive grayscale images (left, centre). Sample image registration of the two frames under translational and rotational transformations using imregister*^†^*(right).

#### Rank-based cell identification and flagging of division events

After locating microchannels in consecutive frames, all cells in the frame of interest *f* are ranked within their respective channels: 1 being closest to the dead-end. The *x*-component of each cell’s centroid is sufficient for ranking due to the orientation of the device. For the first frame in the time series, all cells are given näıve labels with no cells designated as daughters of a division. Cell rank, labelling, and microchannel index were then stored and passed on to the next tracking iteration using the next frame.

For all *f >* 1, a rank-based comparison of cell area between frames *f* and *f −* 1 was performed to detect division events. For two channels paired across consecutive frames, denote *r*_0_ and *r*_1_ as rank-based indices for the cells/ranks of interest in frames *f −* 1 and *f* respectively. Let *A*(*r*_0_) and *A*(*r*_1_) be the areas of the cells of rank *r*_0_ and *r*_1_ in their associated frames. The procedure begins by checking the first cells in the channel: *r*_0_ = *r*_1_ = 1. The proportional change in area *φ_A_* is then calculated:

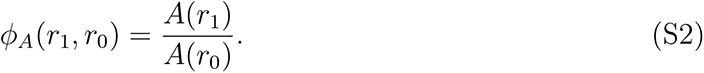

The conditions of the experiments presented here do not anticipate any decreases in cell area unattributable to cell division: under stable nutritional environments, both growth media used are capable of sustaining long-term, balanced growth of all assayed strains. A lower empirical threshold *φ_A_ <* 0.7 was established to mark significant area decreases as cell divisions. Note that since healthy divisions are roughly symmetric in *E. coli*, the next cell along the microchannel should also be relatively small and thus *φ_A_*(*r*_1_ +1*, r*_0_) should also fall below this threshold. The labelling procedure is demonstrated in Fig. S21); the algorithm is explicitly described in bugpipe documentation.

### CHARACTERIZATION OF SEGMENTED CELLS

Fluorescence images and the cell boundaries retrieved from segmentation are used to measure cell size, shape, promoter expression, and the time derivatives of these quantities. As a rod-shaped bacteria, the geometry of an *E. coli* cell can be approximated to a cylinder with hemispherical caps. As shown, the cells grow in the mother machine with their long axis parallel to the plane of observation so that their length and width can be ascertained in addition to their cross sectional area and fluorescence intensity.

The function `getCellPropsMP`, called by `cellTrackerMP`, takes boundary coordinates and a fluorescence intensity matrix as input to determine single cell properties using methods summarized in Table S1 (see also Fig. S22)

Alternative ways to estimate cell volume and the overall geometry of the cell were explored (not shown here); the spherocylindrical model provided the greatest consistency with the volume estimated by the rotation of the boundary contour. During the shift, tapered cells may form due to cell width dynamics [38, 63]. In our experiment, these tapered cells are not easily visible by eye, due to the limited resolution (Fig. S23A). However, we could quantify their contribution by looking at the differences between measured area from segmented perimeter and area computed by width and length measurements assuming a spherocylinder (Fig. S23BC). Quantitatively, this phenomenon is small in our data (about 2%), and does not affect any of our conclusions.

**FIG. S21.**
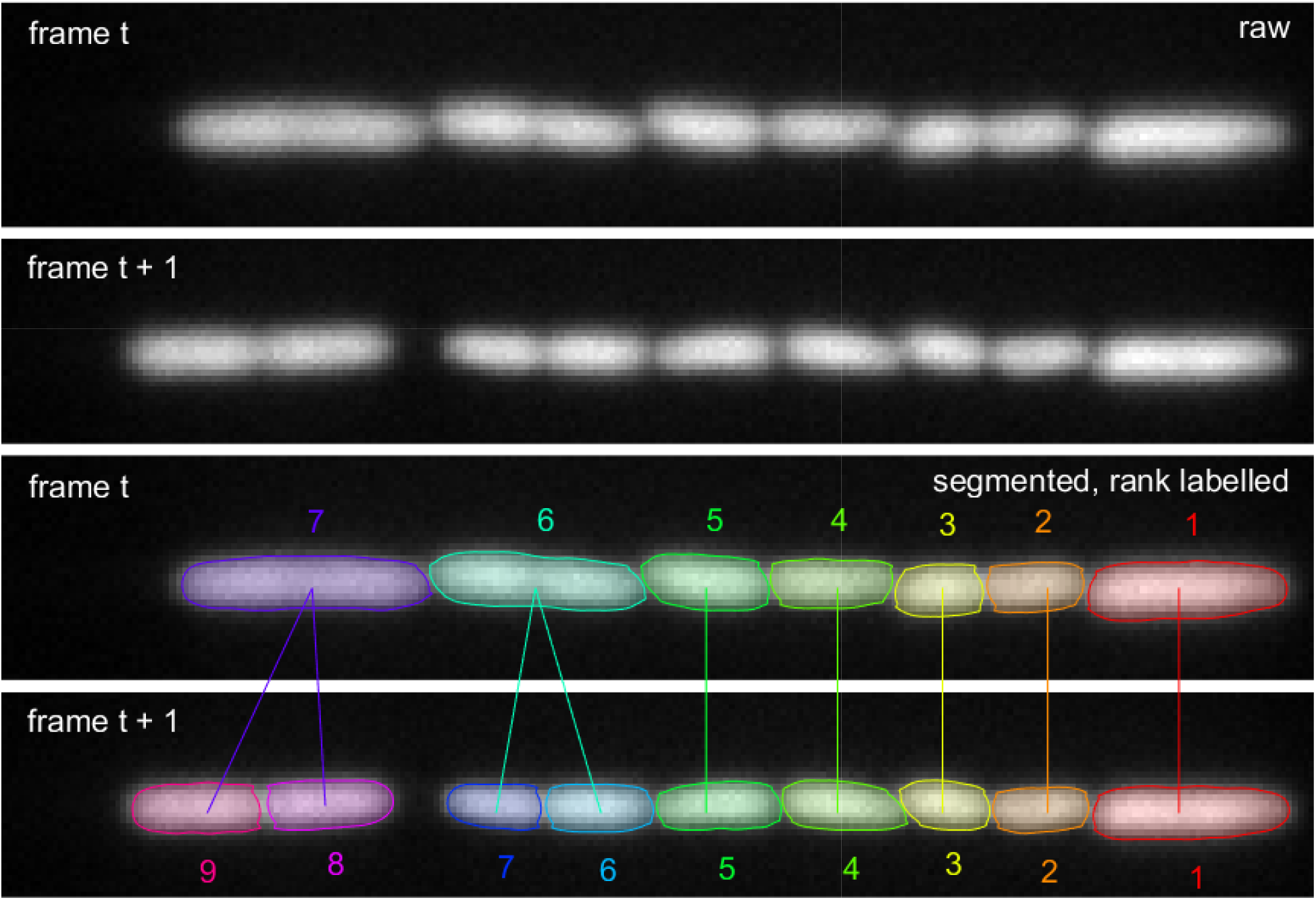
Rank-based comparison of cell area to track cells and division events. A given microchannel is matched between consecutive frames (top two panels). Segmented cells are ranked according to centre of mass distance from the dead-end and changes in area are used to detect division events (bottom two panels). Here, the sixth ranked cell shows a dramatic decrease in area between frames. Finding the area of the next cell in line to be sufficiently small, the procedure determines that a division has indeed occurred and that the sixth and seventh ranked cells in frame *t* + 1 are the daughters of the sixth ranked cell in frame *t*.

### BUGPIPE: SCRIPT IMPLEMENTATION AND DATA MANAGEMENT

Application of the segmentation and tracking procedures is quite straightforward, as demonstrated in the subsection below. There was, however, an additional computational challenge following image processing and analysis: the high throughput and high spatiotemporal resolution nature of acquisition necessitated the development of a fast and flexible means of data manipulation. Custom MATLAB classes were written to handle and analyse these large, non-uniform datasets.

**TABLE S1.** Summary of key single cell measurements

|  |  |  |
| --- | --- | --- |
| area | $A$ | area enclosed by boundary points, including edge pixels |
| length | $L$ | greatest pairwise distance between boundary points along the first principal axis (found by `pca`<sup>†</sup>) |
| width | $w$ | width assuming the projected area is a rectangle with semi-circular caps of radius $\frac{w}{2}$, i.e. $w = \frac{2[L - \sqrt{L^2 - (4 - \pi)A}]}{4 - \pi}$ |
| surface area | $S$ | $S = \pi L w$ |
| volume | $V$ | $V = \frac{4}{3}\pi \left(\frac{w}{2}\right)^3 + \pi \left(\frac{w}{2}\right) (L - w)$ |
| volumetric growth rate | $\alpha_{\text{inst}}$ | instantaneous rate assuming exponential growth: $\alpha_{\text{inst}} = \frac{1}{V} \frac{dV}{dt}$ |
| surface synthesis rate | $\beta$ | instantaneous surface area growth rate assuming it is proportional to volume, i.e. $\beta = \frac{1}{V} \frac{dS}{dt}$ |
| cell cycle growth rate | $\alpha_{\text{cc}}$ | overall volumetric growth rate assuming exponential growth, determined from the slope of the linear fitting of $\log(V)$ vs $t$ for all points of the cell cycle |
| interdivision time | $\tau$ | time between birth and division, estimated as the time between a cell's first and last frames plus the time between consecutive frames |
| promoter/gene expression | $F$ | total fluorescence intensity of the segmented pixels corresponding to $A$ |
| protein concentration | $C$ | concentration of fluorescent protein within the cell: $C = F/V$ |

#### Sample script and directory organization

The following MATLAB script can be taken as a review of the image processing steps described in previous sections and should act as a guide for future users of this package. Note that the script can be easily modified to decrease processing times further by taking advantage of MATLAB’s Parallel Computing Toolbox. Indeed, parallel loops (`spmd`*^†^* and `parfor`*^†^*) were used to process all images pertaining to the data presented in this work.

**FIG. S22.**
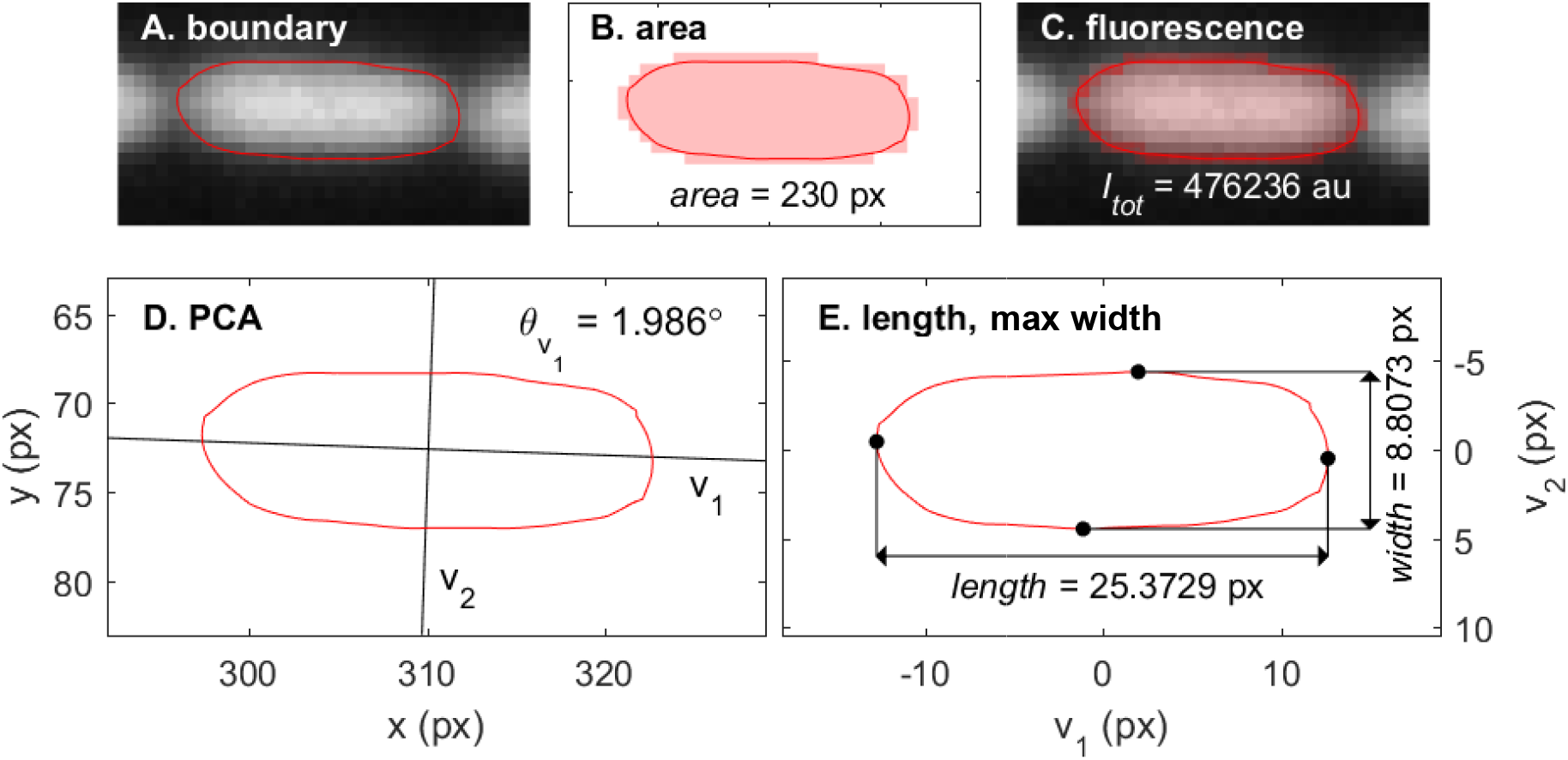
Sample cell measurements retrieved from segmentation of a single cell. **A** Boundary coordinates found overlayed on the original grayscale image. **B** Total cross sectional area calculated as the area enclosed by the boundary, including edge pixels. **C** Total fluorescence intensity associated with the enclosed area. **D** Principal component analysis on boundary to determine axis of orientation. **E** Cell presented in principal axis space, where length is the maximum distance between boundary points along the first axis. Along the second axis, maximum width also shown.

**FIG. S23.**
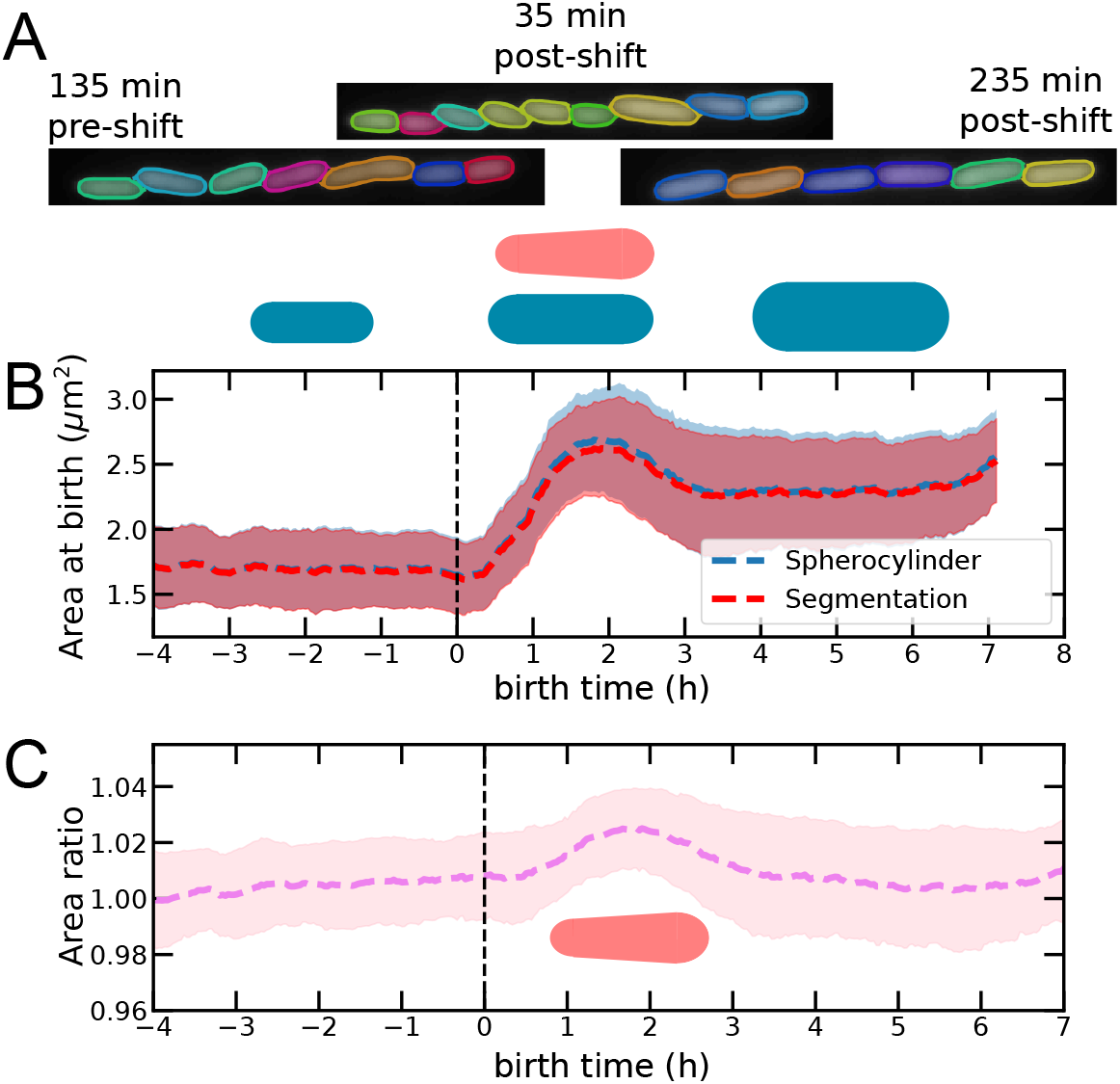
Tapered cells determine minor deviations in estimated volume. **A** Tapered cells are not easily visible by eye at the resolution of our experiments. **B** Comparison of area at birth computed from width and length assuming a spherocylinder, and area at birth determined by perimeter segmentation shows a different during the shift, likely due to the existence of tapered cells [63]. **C** The ratio of assumed to measured area deviates of at most 2% during the shift. The same analysis shows smaller deviations (1-1.5%) if all cell areas are considered and not only area at birth.

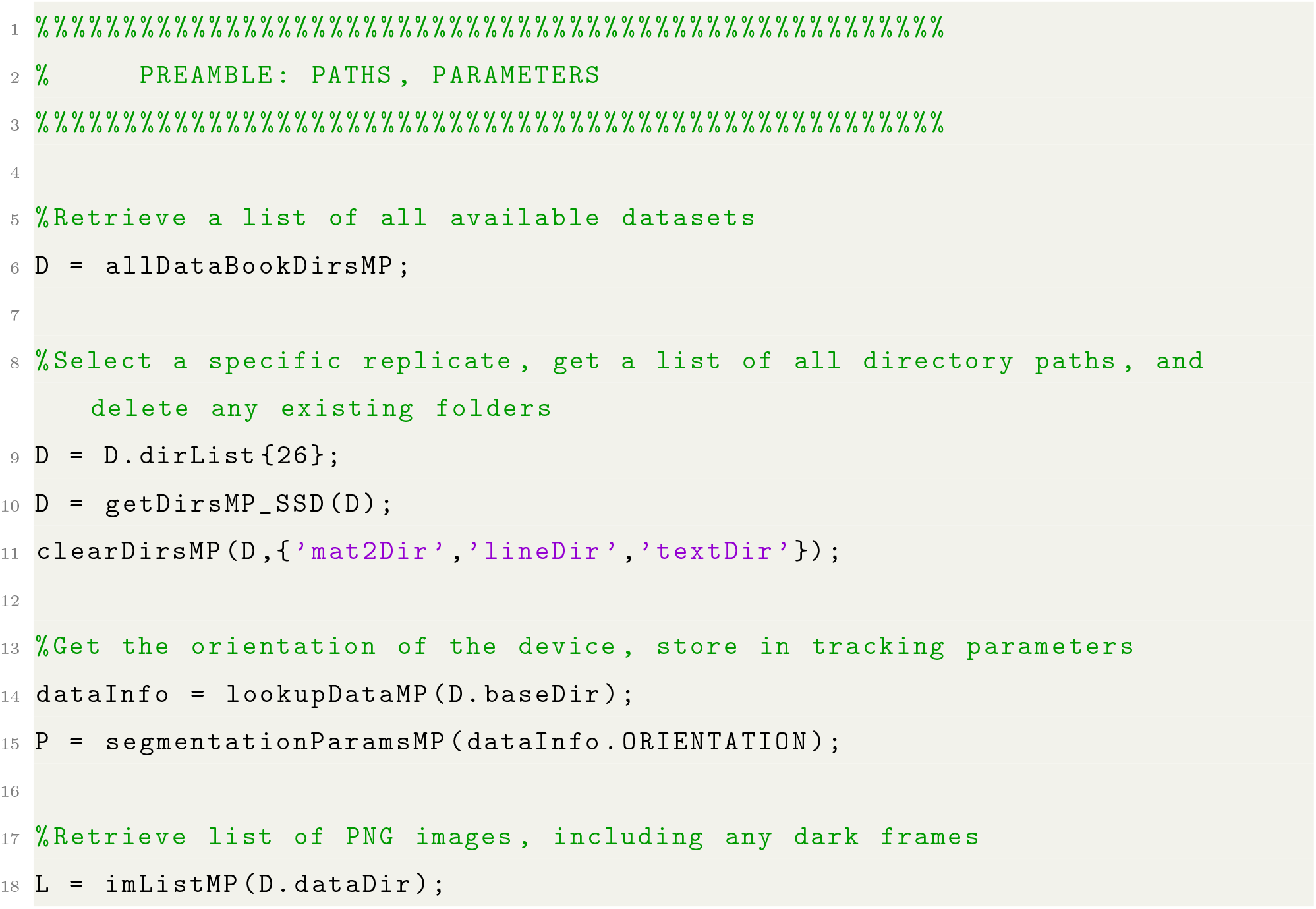
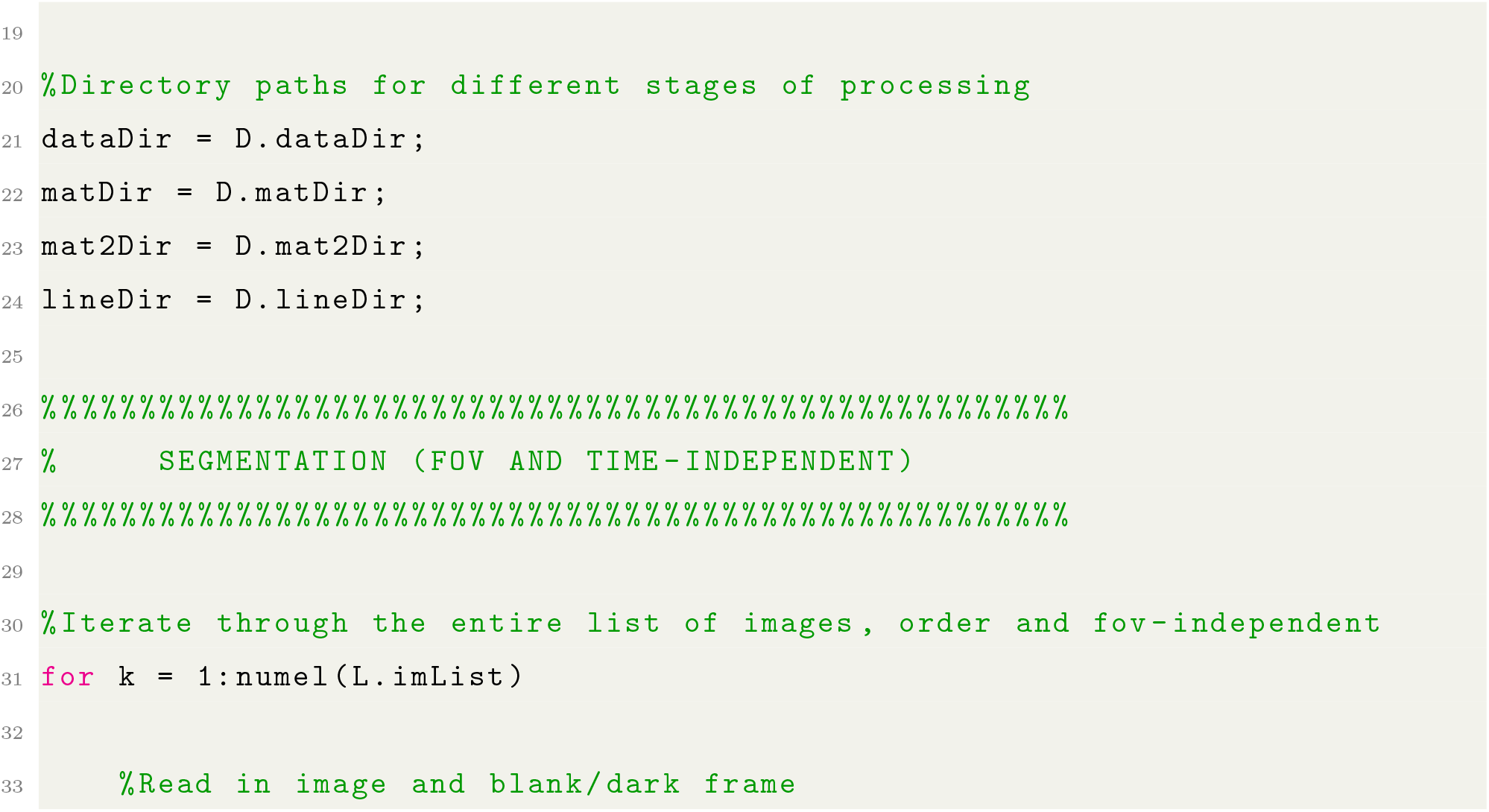
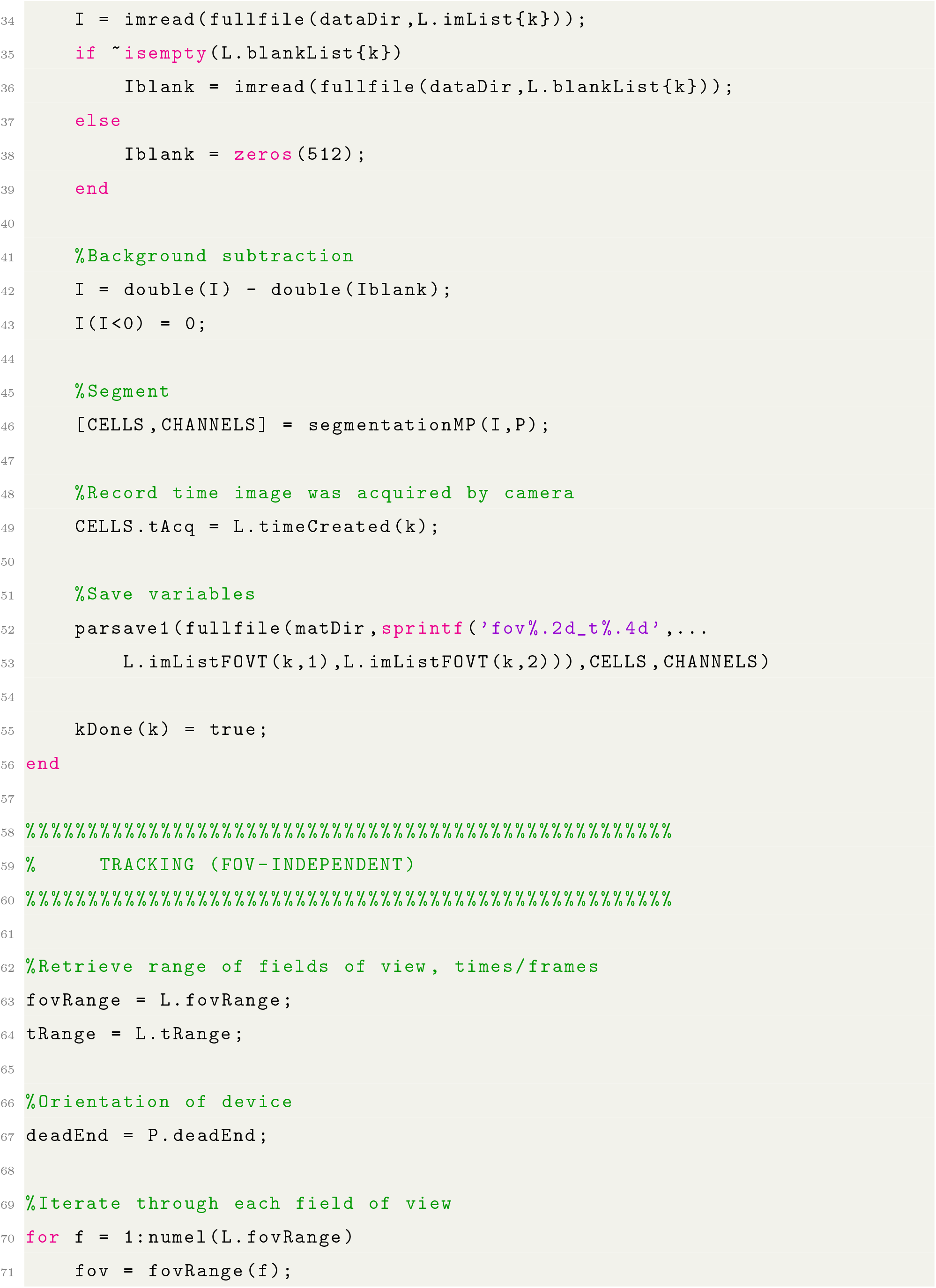
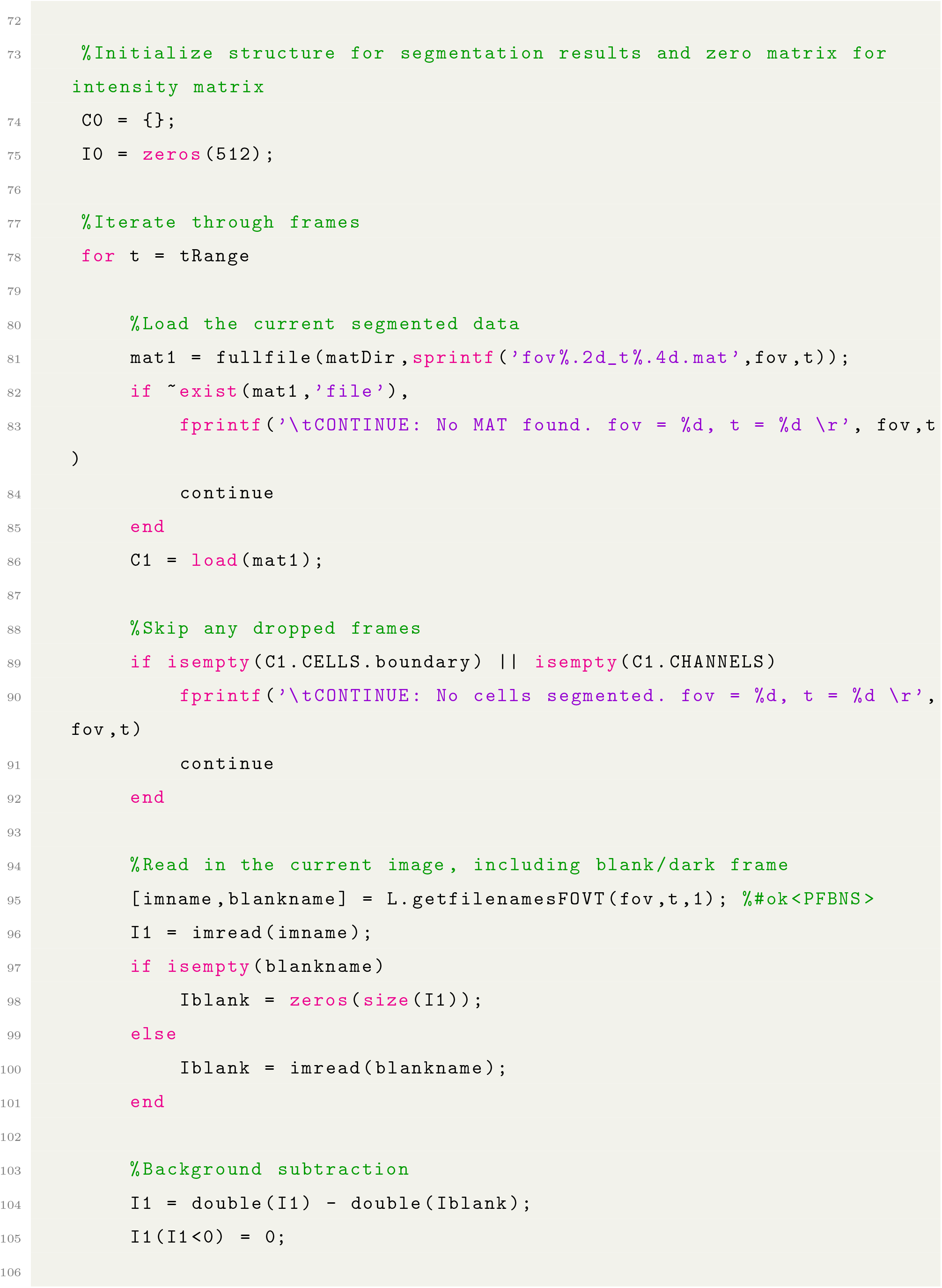
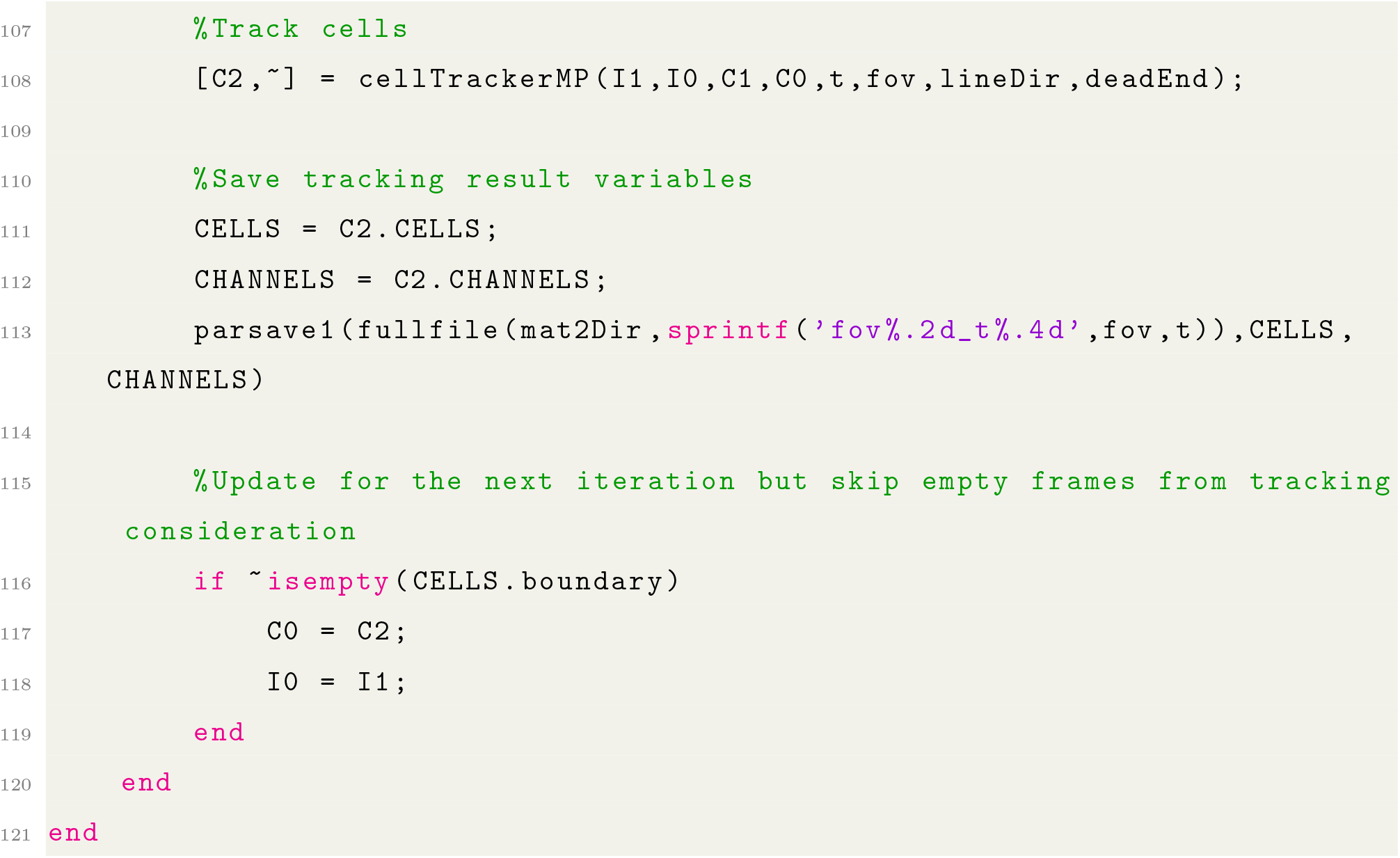

##### Directory structure

Variables are saved at each stage of the procedure, with the final processing step being a conversion from MAT format to text file by the function `mat2txtMP`. The user can delete intermediary variables and directories where needed. A list of the output variables in their associated directories as laid out in the example script are presented in Table S2, with intermediary outputs indicated by (*). More detailed information on each variable is present in the relevant function’s description.

#### Data handling classes

Under the acquisition settings descibed in the main text, about 2,000 cells were imaged every 5 minutes in a typical sample. For the usual 18 h experiment, this translates to over 400,000 cell snapshots and 20,000 fully tracked cell cycles per replicate. Two handling classes were developed to manipulate this data effectively: `dataMP2`, which handles specific replicates; and `dataLabMP`, which manages several replicates at once. These objects allow the user to perform calculations on and retrieve statistics from different populations: whether it be of single cell cycles, specific lineages, whole replicates, or multiple datasets. The complete documentation, including example analyses, can be found on https://github.com/panlilio/bugpipe.

**TABLE S2.**
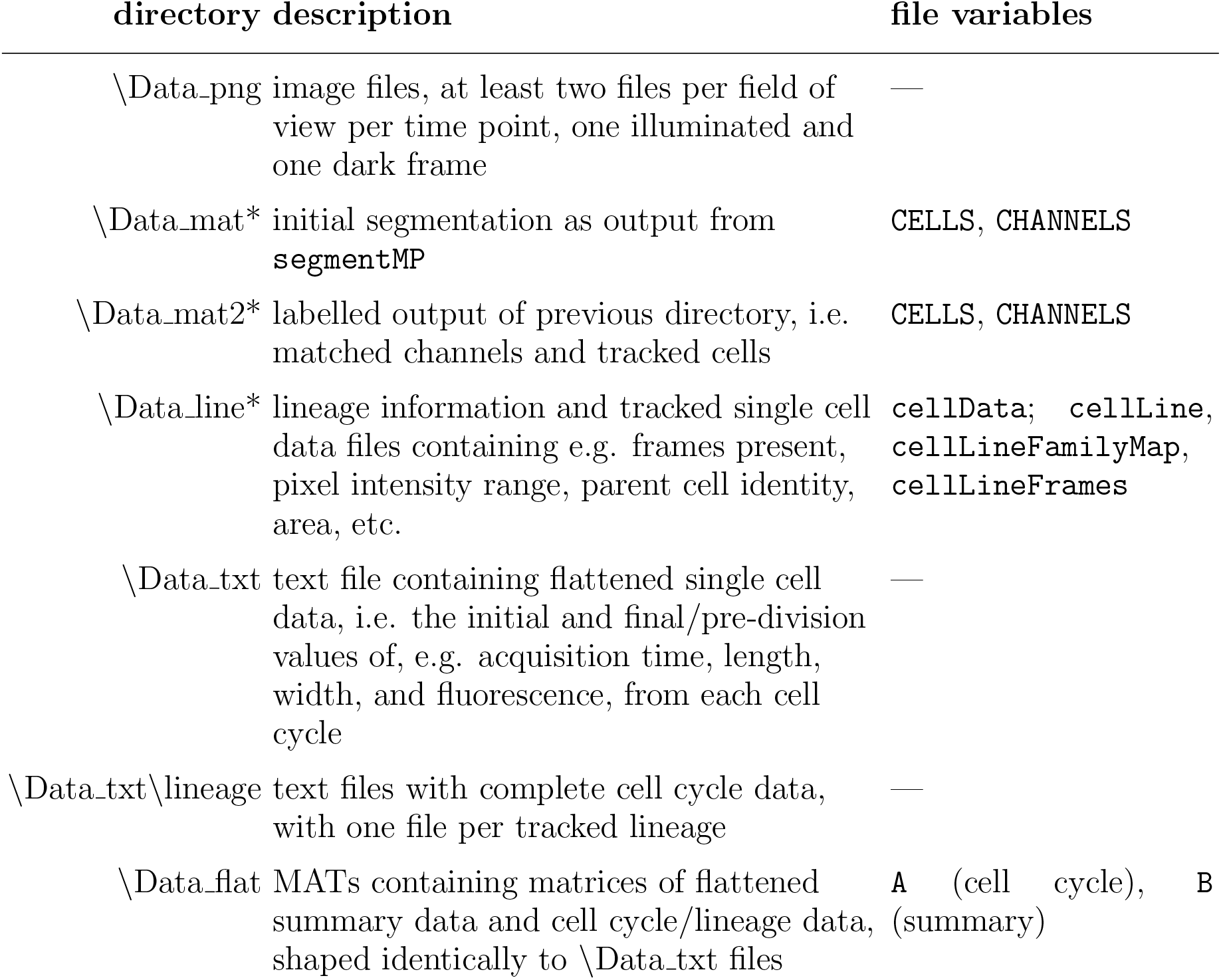
Organizational structure of segmentation results

| directory | description | file variables |
| --- | --- | --- |
| \Data_png | image files, at least two files per field of view per time point, one illuminated and one dark frame | — |
| \Data_mat* | initial segmentation as output from `segmentMP` | CELLS, CHANNELS |
| \Data_mat2* | labelled output of previous directory, i.e. matched channels and tracked cells | CELLS, CHANNELS |
| \Data_line* | lineage information and tracked single cell data files containing e.g. frames present, pixel intensity range, parent cell identity, area, etc. | `cellData`; `cellLine`,<br>`cellLineFamilyMap`,<br>`cellLineFrames` |
| \Data_txt | text file containing flattened single cell data, i.e. the initial and final/pre-division values of, e.g. acquisition time, length, width, and fluorescence, from each cell cycle | — |
| \Data_txt\lineage | text files with complete cell cycle data, with one file per tracked lineage | — |
| \Data_flat | MATs containing matrices of flattened summary data and cell cycle/lineage data, shaped identically to \Data_txt files | A (cell cycle), B (summary) |

#### Applied data filters

Following the highly automated cell segmentation and tracking procedures, there were a few common data filters applied for subsequent analyses. Specifically, we required that only cells meeting the following criteria were considered:

1. The entire life cycle was observed, i.e. both parent and daughter(s) were at least partially tracked so that a division event was flagged both directly before and after the cell cycle.
2. The division event terminating the cell cycle of interest must split the mother into near-symmetrical daughters, specifically each daughter occupying 40-60% of the mother’s final cross sectional area. This step aims to eliminate filamentous cells and segmentation artifacts.
3. The observed exponential growth rate corresponds to a doubling time that is greater than 10 minutes but finite. Note that the physiological lower limit is 15-20 min.

Note that the last filter is largely unnecessary for the vast majority of datasets presented in this work. For example, in replicate 20160325, less than 0.3% (104 out of 37,570) of segmented cells lie outside of this reasonable growth rate interval.

